# Explaining plant trait variation in response to soil water availability using an optimal height-growth model

**DOI:** 10.1101/2024.01.23.576942

**Authors:** Isaac R. Towers, Andrew O’Reilly-Nugent, Manon E.B. Sabot, Peter A. Vesk, Daniel S. Falster

**Affiliations:** The School of Biological, Earth and Environmental Sciences, The University of New South Wales, NSW, 2052, Australia; Climate Friendly, Sydney, NSW, 2052, Australia; Max Planck Insitute for Biogeochemistry, 07745 Jena, Germany; ARC Centre of Excellence for Climate Extremes and Climate Change Research Centre, The University of New South Wales, NSW, 2052, Australia; School of Agriculture, Food and Ecosystem Sciences, The University of Melbourne, Parkville, VIC, 3010, Australia

**Keywords:** leaf mass per area, huber value, wood density, sapwood-specific conductivity, eco-evolutionary optimality, stomata, hydraulics, trait change

## Abstract

Climate change is expected to bring about changes in precipitation and temperature regimes that, together with rising atmospheric CO_2_ concentrations, will likely reorganise the functional trait composition of ecosystems. Predicting plant trait responses to emerging environmental conditions including, in particular, water availability, is a tremendous challenge, but is one that eco-evolutionary optimality theory (EEO) can help us undertake. However, most EEO approaches are based on the hypothesis that traits are selected to maximise carbon assimilation which omits the important role that size growth plays in determining fitness outcomes. Using a height-growth based EEO framework, we predict magnitude and directional shifts in four key traits: leaf mass per area, sapwood area to leaf area ratio (Huber value), wood density and sapwood-specific conductivity in response to variation in soil moisture availability, atmospheric aridity, CO_2_ and light availability. Consistent with empirical patterns, we predict that trait optima shift from resource-acquisitive strategies characterised by low tissue constructions costs and high rates of tissue turnover and sapwood conductivity to resource-conservative strategies - characterised by low rates of tissue turnover and greater xylem embolism resistance - as conditions become increasingly dry. The EEO model that we use here highlights the important role that both carbon assimilation and tissue construction costs jointly play in predicting the response of trait optima to the environment, laying the groundwork for future height-growth based EEO models aiming to predict shifts in the functional composition of ecosystems in response to global change.

## 2 Introduction

The extraordinary diversity of plant species on Earth is testament to the existence of trade-offs in the ecophysiological strategies employed by plants to grow, survive and reproduce. Biophysical constraints on plant phenotype mean that the benefits conferred by a given strategy also impose costs to fitness such that no organism is perfectly adapted to all conditions (Laughlin 2018). As such, we expect to observe shifts in plant community composition as conditions change across space and time. Precipitation, soil types, and thus soil moisture, vary widely across the globe (Fick & Hijmans 2017), and there is ample empirical evidence that plants exhibit strong responses in their traits to water availability. These responses concern traits related to hydraulic function (such as sapwood area and conductivity), tissue construction (such as leaf mass per area and wood density) and life cycle (maximum height) (Niinemets 2001; Wright *et al*. 2004; Moles *et al*. 2009; Towers *et al*. 2023). Although we understand the mechanisms underlying some of these patterns, more theory is needed to explain the selective forces causing these patterns to emerge across large spatial scales and at what temporal scales they occur.

The need for further theoretical understanding is motivated by the rapid environmental changes that Earth is experiencing. In addition to rising temperature (and thus), atmospheric dryness and carbon dioxide concentrations, climate change is expected to bring about changes in precipitation regimes across the globe (IPCC 2023), and thereby modifying the functional composition of ecosystems as populations and species adapt, migrate, or are driven extinct under new climatic conditions (Zhu *et al*. 2012; Crimmins *et al*. 2011). Indeed, there is already evidence showing an increase in the predominance of drought-affiliated species in ecosystems that are becoming drier and more seasonal and, on the other hand, the invasion of mesophilic species in locations that are becoming wetter (Feeley *et al*. 2011, 2020; Fauset *et al*. 2012). Without theoretical predictions on how these changes may determine the favourability of different plant strategies, we are unable to anticipate the direction and rate of change in the composition of future vegetation.

Eco-evolutionary optimality frameworks (EEO) offers a means to develop process-based hypotheses for how and why traits respond to the environment (Harrison *et al*. 2021). Under EEO, environmentally-driven trait patterns are hypothesised to emerge from a selection of trait combinations that maximises reproductive fitness. However, due to the difficulty of estimating reproductive fitness, EEO-based models typically maximise some other variable as a tractable proxy. For example, many existing implementations of EEO optimise carbon assimilation on an instantaneous-basis or through time at the leaf, plant or ecosystem-scale. These models have been shown to successfully replicate observed environmental responses for a number of traits and variables including, among others, the intercellular to atmospheric ratio of CO_2_ (Wang *et al*. 2017), leaf maximum photosynthetic carboxylation capacity (Dong *et al*. 2017) and the sapwood to leaf area ratio (Xu *et al*. 2021; Trugman *et al*. 2019). Despite success, it has been argued that optimality approaches that focus on carbon acquisition alone limit the range of traits that can be represented by EEO because traits can also affect the construction cost of and relative allocation towards different plant tissues instead of, or in addition to, carbon uptake (Falster *et al*. 2018; Bartlett *et al*. 2019; Dong *et al*. 2022). In such cases, trait-environment gradients emerging from selection may only become evident when considering how carbon production translates into size growth but, because there is little consensus on how to achieve this translation, it has seldom been investigated (but see Potkay & Feng 2023).

Trait growth theory (TGT) integrates the effects of traits on both biomass production and plant construction costs on growth rates (Falster *et al*. 2011, 2018; Gibert *et al*. 2016; Westoby *et al*. 2022). This is achieved by linking both production and construction to measurable traits and the tradeoffs encoded by these traits, to describe how additional biomass is allocated to different tissues as the plant grows. Thus far, TGT has been used to provide mechanistic explanations for a variety of empirical phenomena including the hump-shaped change in height growth rate of biomass with size, the size-dependent effect of traits on growth, and effect of traits on shade tolerance (Falster *et al*. 2017). The system of equations that make up TGT mean that, in principle, the model can be extended to include the effect of any abiotic factor on growth, so long as it influences either biomass production and/or allocation. The flexibility of TGT is particularly suited for explaining empirical patterns in the occurrence of traits along environmental gradients.

Recent advances in the representation of trade-offs between photosynthetic and hydraulic functions offer a mechanistic, first-principles pathway to link stomatal response and traits to drying soil (Wolf *et al*. 2016). As soils dry, plants face an unavoidable trade-off between keeping their stomata open to maintain photosynthesis and closing their stomata to minimise drought-induced damaged to the water transport pathway. In contrast to empirical stomatal models, a number of optimal stomatal models assume plants to be efficient water users (Cowan & Farquhar 1977) that actively regulate the benefits of carbon acquisition relative to the costs of water loss. Beyond their demonstrated performance in predicting observed stomatal conductance (Sabot *et al*. 2022), optimal stomatal models have the added advantage of being parameterised with measurable hydraulic traits, thereby mechanistically linking traits to stomatal behaviour and thus, to fitness.

Here, we integrate an optimal stomatal behaviour model into the TGT framework to generate predictions about how traits should respond to gradients in soil moisture, atmospheric vapour pressure deficit, carbon dioxide and light availability. The stomatal behaviour model is based on maximising carbon acquisition after accounting for costs associated with water acquisition. We use plant growth rate as a proxy for fitness, thereby integrating effects of traits on biomass production and plant construction. Our primary goal is to qualitatively capture the directions of empirically observed trait responses to changes in soil water availability as an emergent outcome of an EEO model based on height growth rates. We then investigate whether traits are predicted to also shift with plant size, either through influencing the relative allocation of carbon to different tissues or the hydraulic transport pathway, as shifts in traits with size are also widely observed in nature (Table 1). Finally, by extending our approach to optimise across multiple trait dimensions simultaneously, we investigate our framework’s ability to explain species turnover across soil moisture gradients in terms of fitness (as represented by height-growth rate).

**Table 1:** Phenomena analysed in the model and empirical evidence for phenomena. Arrows shows the direction of the empirical trait response

| Phenomenon | Response of trait |  |  |  |
| --- | --- | --- | --- | --- |
| | $\phi$ | $\rho$ | $K_{s,max}$ | $\theta$ |
| **Variation in trait across environments** |  |  |  |  |
| With soil dryness | $\uparrow^1$ | $\uparrow^{1,2,3}$ | $\downarrow^{4,5}$ | $\uparrow^1$ |
| With atmospheric vapour pressure deficit | $\uparrow^6$ | $\uparrow^7$ | $\uparrow^{8,9}$ | $\uparrow^{10}$ |
| With atmospheric carbon dioxide concentration | $\uparrow^{11,12,13}$ | $\downarrow^{14}$ | $\downarrow^{14,15}$ | $\downarrow^{14}$ |
| With light availability | $\uparrow^{13,16,17}$ | $\downarrow^{18}$ | $\uparrow^{19}$ | $\uparrow^{20}$ |
| **Variation in trait within individuals** |  |  |  |  |
| Change with ↑ size | $\uparrow^{21}$ | $\uparrow^{22}$ | $\uparrow^{23}$ | $\uparrow^{24}$ |

<sup>1</sup> Towers *et al.* (2023) <sup>2</sup> Onoda *et al.* (2010) <sup>3</sup> Pickup *et al.* (2005) <sup>4</sup> Gleason *et al.* (2013) <sup>5</sup> Tavares *et al.* (2023) <sup>6</sup> Westerband *et al.* (2023) <sup>7</sup> Rocha *et al.* (2020) <sup>8</sup> Liu *et al.* (2021) <sup>9</sup> Olson *et al.* (2020) <sup>10</sup> Mencuccini & Grace (1995) <sup>11</sup> Hikosaka *et al.* (2005) <sup>12</sup> Pritchard *et al.* (1999) <sup>13</sup> Poorter *et al.* (2009) <sup>14</sup> Bobich *et al.* (2010) <sup>15</sup> Eguchi *et al.* (2008) <sup>16</sup> Ellsworth & Reich (1992) <sup>17</sup> Neyret *et al.* (2016) <sup>18</sup> Poorter *et al.* (2019) <sup>19</sup> Sellin *et al.* (2010) <sup>20</sup> de Oliveira *et al.* (2023) <sup>21</sup> Westoby *et al.* (2022) <sup>22</sup> Hietz *et al.* (2013) <sup>23</sup> Rungwattana & Hietz (2018) <sup>24</sup> McDowell *et al.* (2002)

## 3 Methods

### 3.1 Trait growth theory

TGT describes how traits influence plant growth rates and how this influence changes with plant size and the environment (Falster *et al*. 2018). The framework considers how whole-plant carbon acquisition translates into the growth of different plant tissues, each of which are tracked explicitly and are related via allometric assumptions of plant construction. In this analysis, we focus on the height growth rate of an individual plant as a proxy for plant fitness in a given environment. As we do not consider differences in canopy architecture amongst plants, the height growth rate is directly related to leaf area growth rate and the two can be used interchangeably. Indeed, size growth (in either leaf area or height) is arguably the primary goal of a young plant aiming to develop a canopy and reproduce.

According to TGT, the height growth rate of an individual, 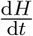, potentially varies with traits (here denoted by the letter *x*), size (*H*) and environment (*E*), as the outcome of four processes:

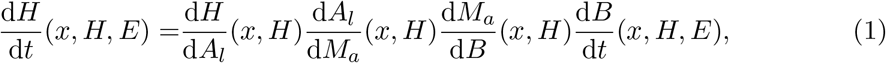

where 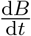 describes the growth of vegetative biomass, 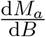 describes the fraction of biomass available for allocation to growth of living tissues (*M*_*a*_) after allocation to reproduction; 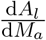 describes the change in canopy leaf area (*A*_*l*_) for a given unit of biomass allocated to growth; and 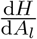 describes the allometric relationship between plant height and crown size. For the present study, we simplify the analysis by assuming that plants do not invest in reproduction and that 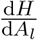 is invariant across taxa, which is reasonable for woody, tree-like taxa (Falster *et al*. 2015). Thus, in our analysis, variation in 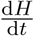 is realised only through variation in 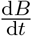 and the relative allocation of mass to different tissues 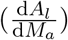.

The term 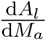 can be further decomposed into the marginal increase in mass of each plant component required to support an additional unit of leaf area:

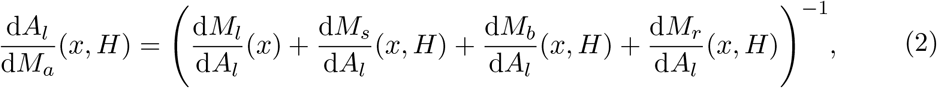

where *M*_*s*_, *M*_*b*_ and *M*_*r*_ are the sapwood, bark and root mass, respectively. Tissue mass is linked to leaf area according to a series of functional-balance equations which describe the relationship between plant properties (Falster *et al*. 2018). Here, 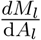 is the leaf mass per area, (*ϕ*), while 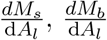 and 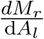 emerge from the pipe model which assumes that sapwood, bark and root surface areas scale proportionally with leaf area (Shinozaki *et al*.1964). Traits such as leaf mass per area can influence 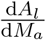 in the model by moderating the area by mass ratio of a given plant tissue or the ratio of leaf area to the area of other tissues.

Biomass growth is modelled as the net increase in biomass resulting from photosynthesis after accounting for losses related to damage to the hydraulic pathway incurred by transpiration, respiration and turnover of tissues:

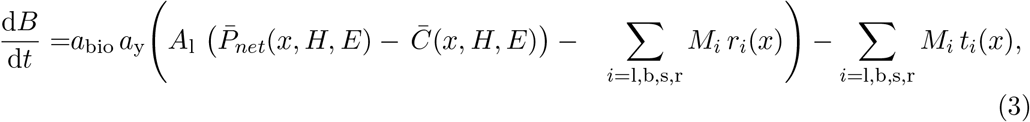

whereby the whole-canopy rates of net photosynthesis and hydraulic cost are found by multiplying the total photosynthetic surface *A*_*l*_ by the average rate of leaf-level net photosynthesis, 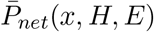 and leaf-level hydraulic cost, 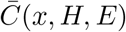, respectively, which are themselves dependent on traits, plant size, the and environment (see Section 3.2). Tissue-specific rates of respiration, *r*_*i*_ and turnover *t*_*i*_ are mass-based and are calculated by summing across the mass of each plant tissue (*i*). *a*_y_ and *a*_bio_ are constants being the fraction of carbon per unit of biomass and the conversion rate between CO_2_ and biomass, respectively (Table 2).

**Table 2:**
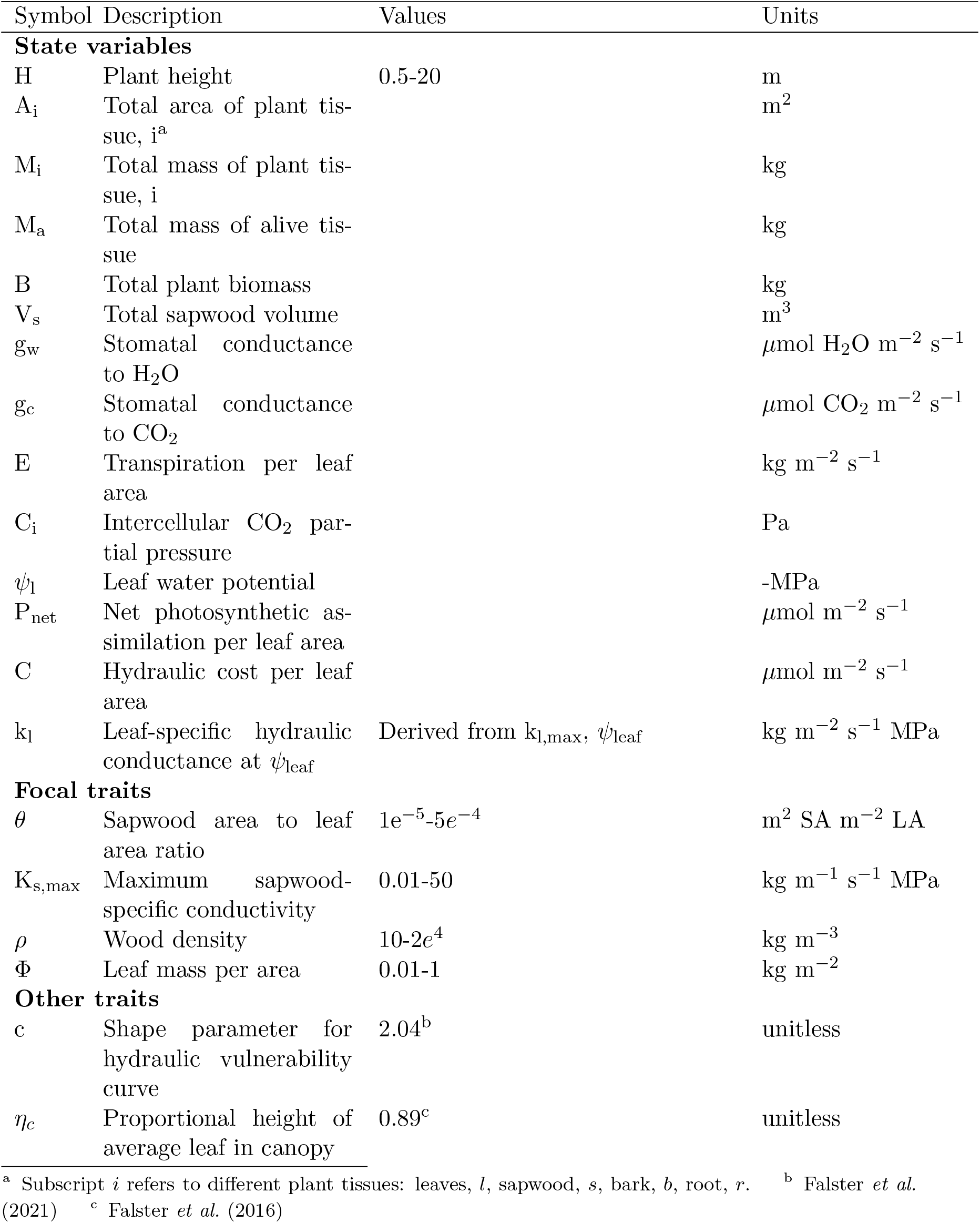
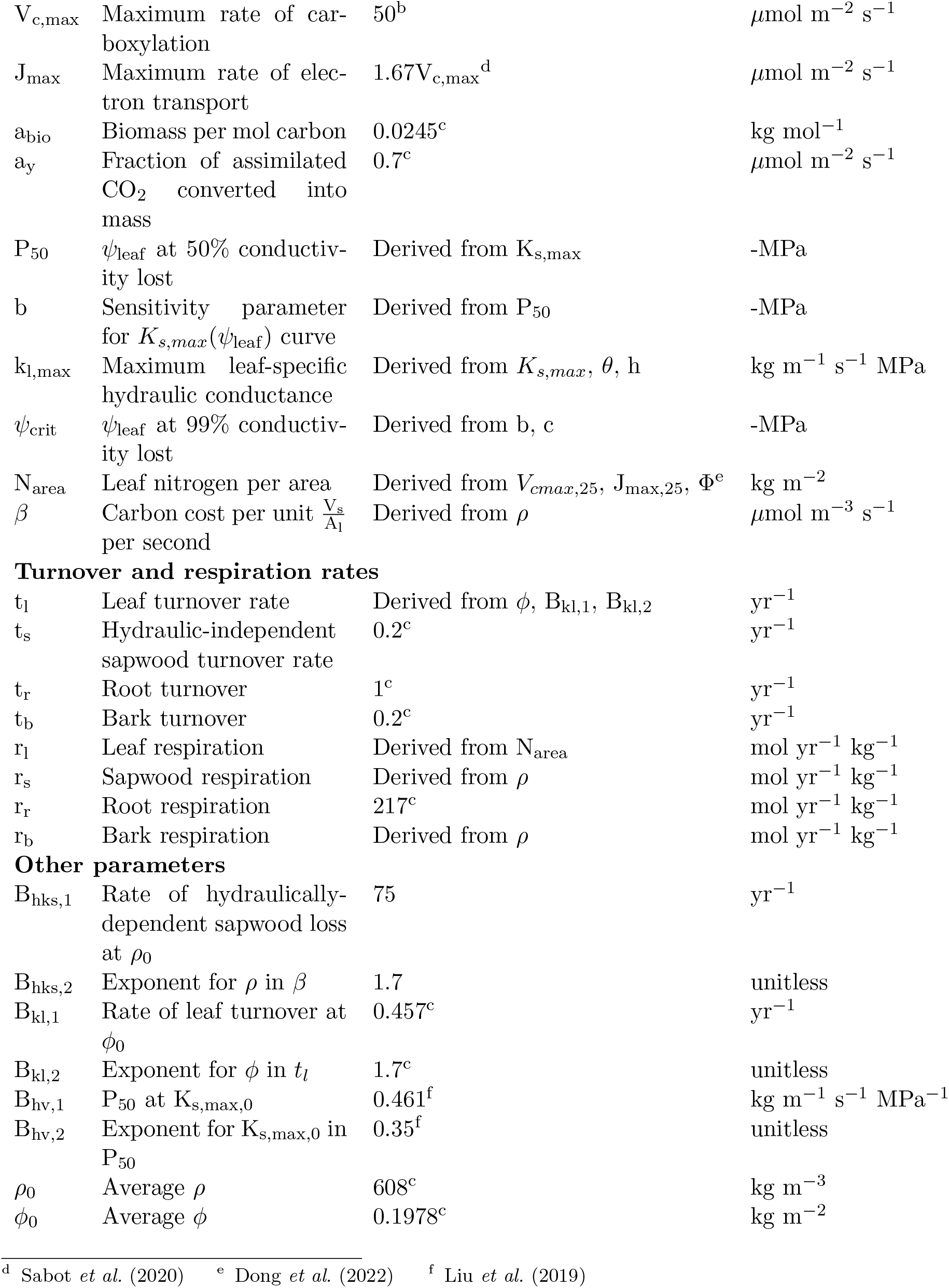
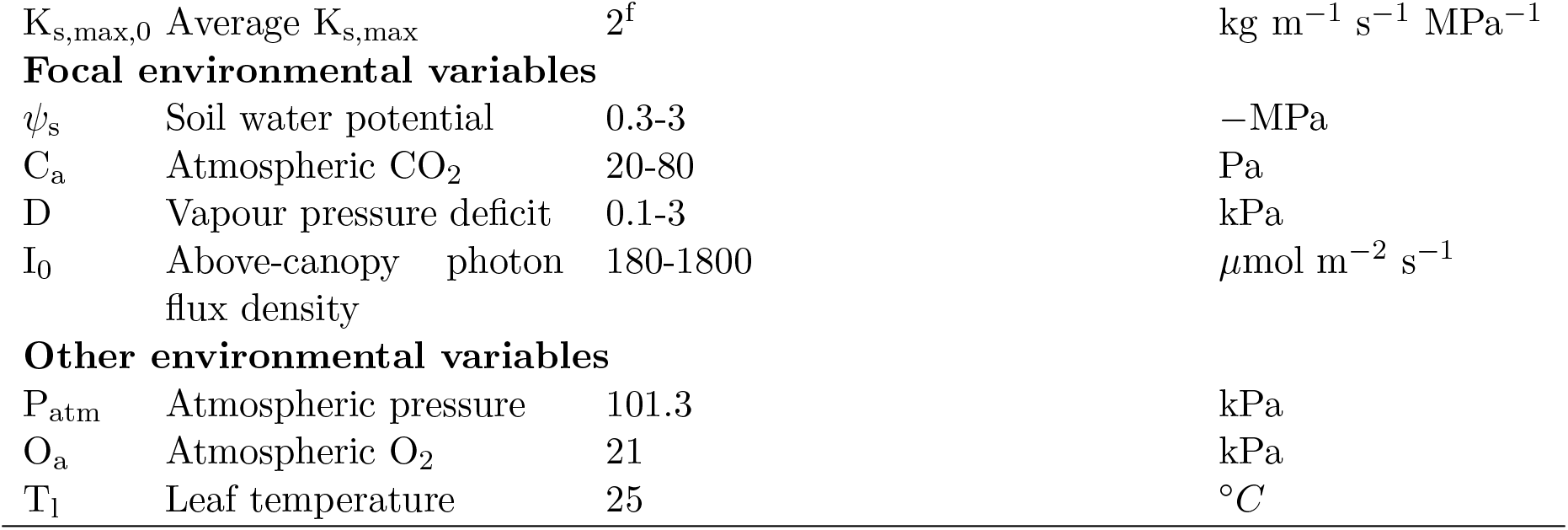
Variable descriptions, tested parameter values and units. Values for state variables left intentionally blank as these vary dynamically.

| Symbol | Description | Values | Units |
| --- | --- | --- | --- |
| **State variables** |  |  |  |
| H | Plant height | 0.5-20 | m |
| A<sub>i</sub> | Total area of plant tissue, i<sup>a</sup> |  | m<sup>2</sup> |
| M<sub>i</sub> | Total mass of plant tissue, i |  | kg |
| M<sub>a</sub> | Total mass of alive tissue |  | kg |
| B | Total plant biomass |  | kg |
| V<sub>s</sub> | Total sapwood volume |  | m<sup>3</sup> |
| g<sub>w</sub> | Stomatal conductance to H<sub>2</sub>O |  | μmol H<sub>2</sub>O m<sup>-2</sup> s<sup>-1</sup> |
| g<sub>c</sub> | Stomatal conductance to CO<sub>2</sub> |  | μmol CO<sub>2</sub> m<sup>-2</sup> s<sup>-1</sup> |
| E | Transpiration per leaf area |  | kg m<sup>-2</sup> s<sup>-1</sup> |
| C<sub>i</sub> | Intercellular CO<sub>2</sub> partial pressure |  | Pa |
| ψ<sub>l</sub> | Leaf water potential |  | -MPa |
| P<sub>net</sub> | Net photosynthetic assimilation per leaf area |  | μmol m<sup>-2</sup> s<sup>-1</sup> |
| C | Hydraulic cost per leaf area |  | μmol m<sup>-2</sup> s<sup>-1</sup> |
| k<sub>l</sub> | Leaf-specific hydraulic conductance at ψ<sub>leaf</sub> | Derived from k<sub>l,max</sub>, ψ<sub>leaf</sub> | kg m<sup>-2</sup> s<sup>-1</sup> MPa |
| **Focal traits** |  |  |  |
| θ | Sapwood area to leaf area ratio | 1e<sup>-5</sup>-5e<sup>-4</sup> | m<sup>2</sup> SA m<sup>-2</sup> LA |
| K<sub>s,max</sub> | Maximum sapwood-specific conductivity | 0.01-50 | kg m<sup>-1</sup> s<sup>-1</sup> MPa |
| ρ | Wood density | 10-2e<sup>4</sup> | kg m<sup>-3</sup> |
| Φ | Leaf mass per area | 0.01-1 | kg m<sup>-2</sup> |
| **Other traits** |  |  |  |
| c | Shape parameter for hydraulic vulnerability curve | 2.04<sup>b</sup> | unitless |
| η<sub>c</sub> | Proportional height of average leaf in canopy | 0.89<sup>c</sup> | unitless |

<sup>a</sup> Subscript *i* refers to different plant tissues: leaves, *l*, sapwood, *s*, bark, *b*, root, *r*. <sup>b</sup> Falster *et al.* (2021) <sup>c</sup> Falster *et al.* (2016)

|  |  |  |  |
| --- | --- | --- | --- |
| $V_{c,max}$ | Maximum rate of carboxylation | 50<sup>b</sup> | $\mu\text{mol m}^{-2} \text{ s}^{-1}$ |
| $J_{max}$ | Maximum rate of electron transport | $1.67V_{c,max}$<sup>d</sup> | $\mu\text{mol m}^{-2} \text{ s}^{-1}$ |
| $a_{bio}$ | Biomass per mol carbon | 0.0245<sup>c</sup> | $\text{kg mol}^{-1}$ |
| $a_y$ | Fraction of assimilated $\text{CO}_2$ converted into mass | 0.7<sup>c</sup> | $\mu\text{mol m}^{-2} \text{ s}^{-1}$ |
| $P_{50}$ | $\psi_{leaf}$ at 50% conductivity lost | Derived from $K_{s,max}$ | -MPa |
| $b$ | Sensitivity parameter for $K_{s,max}(\psi_{leaf})$ curve | Derived from $P_{50}$ | -MPa |
| $k_{l,max}$ | Maximum leaf-specific hydraulic conductance | Derived from $K_{s,max}$, $\theta$, $h$ | $\text{kg m}^{-1} \text{ s}^{-1} \text{ MPa}$ |
| $\psi_{crit}$ | $\psi_{leaf}$ at 99% conductivity lost | Derived from $b$, $c$ | -MPa |
| $N_{area}$ | Leaf nitrogen per area | Derived from $V_{c,max,25}$, $J_{max,25}$, Φ<sup>e</sup> | $\text{kg m}^{-2}$ |
| $\beta$ | Carbon cost per unit $\frac{V_s}{A_l}$ per second | Derived from $\rho$ | $\mu\text{mol m}^{-3} \text{ s}^{-1}$ |
| Turnover and respiration rates |  |  |  |
| $t_l$ | Leaf turnover rate | Derived from $\phi$, $B_{kl,1}$, $B_{kl,2}$ | $\text{yr}^{-1}$ |
| $t_s$ | Hydraulic-independent sapwood turnover rate | 0.2<sup>c</sup> | $\text{yr}^{-1}$ |
| $t_r$ | Root turnover | 1<sup>c</sup> | $\text{yr}^{-1}$ |
| $t_b$ | Bark turnover | 0.2<sup>c</sup> | $\text{yr}^{-1}$ |
| $r_l$ | Leaf respiration | Derived from $N_{area}$ | $\text{mol yr}^{-1} \text{ kg}^{-1}$ |
| $r_s$ | Sapwood respiration | Derived from $\rho$ | $\text{mol yr}^{-1} \text{ kg}^{-1}$ |
| $r_r$ | Root respiration | 217<sup>c</sup> | $\text{mol yr}^{-1} \text{ kg}^{-1}$ |
| $r_b$ | Bark respiration | Derived from $\rho$ | $\text{mol yr}^{-1} \text{ kg}^{-1}$ |
| Other parameters |  |  |  |
| $B_{hks,1}$ | Rate of hydraulically-dependent sapwood loss at $\rho_0$ | 75 | $\text{yr}^{-1}$ |
| $B_{hks,2}$ | Exponent for $\rho$ in $\beta$ | 1.7 | unitless |
| $B_{kl,1}$ | Rate of leaf turnover at $\phi_0$ | 0.457<sup>c</sup> | $\text{yr}^{-1}$ |
| $B_{kl,2}$ | Exponent for $\phi$ in $t_l$ | 1.7<sup>c</sup> | unitless |
| $B_{hv,1}$ | $P_{50}$ at $K_{s,max,0}$ | 0.461<sup>f</sup> | $\text{kg m}^{-1} \text{ s}^{-1} \text{ MPa}^{-1}$ |
| $B_{hv,2}$ | Exponent for $K_{s,max,0}$ in $P_{50}$ | 0.35<sup>f</sup> | unitless |
| $\rho_0$ | Average $\rho$ | 608<sup>c</sup> | $\text{kg m}^{-3}$ |
| $\phi_0$ | Average $\phi$ | 0.1978<sup>c</sup> | $\text{kg m}^{-2}$ |

<sup>d</sup> Sabot *et al.* (2020) <sup>e</sup> Dong *et al.* (2022) <sup>f</sup> Liu *et al.* (2019)

|  |  |  |  |
| --- | --- | --- | --- |
| $K_{s,max,0}$ | Average $K_{s,max}$ | $2^f$ | $\text{kg m}^{-1} \text{s}^{-1} \text{MPa}^{-1}$ |
| **Focal environmental variables** |  |  |  |
| $\psi_s$ | Soil water potential | 0.3-3 | –MPa |
| $C_a$ | Atmospheric $\text{CO}_2$ | 20-80 | Pa |
| $D$ | Vapour pressure deficit | 0.1-3 | kPa |
| $I_0$ | Above-canopy photon flux density | 180-1800 | $\mu\text{mol m}^{-2} \text{s}^{-1}$ |
| **Other environmental variables** |  |  |  |
| $P_{\text{atm}}$ | Atmospheric pressure | 101.3 | kPa |
| $O_a$ | Atmospheric $\text{O}_2$ | 21 | kPa |
| $T_l$ | Leaf temperature | 25 | $^{\circ}\text{C}$ |

### 3.2 Stomatal behaviour model

We extended the photosynthesis sub-model used in previous implementations of the **plant** model, by incorporating stomatal behavioural responses to the environment into the model. Specifically, inspired by the work of Bartlett *et al*. (2019) linking stomatal behaviour to evolutionary traits on the basis of maximising carbon acquisition relative to hydraulic costs, and following the idea that stomatal optimality should apply at any given instant (Wolf *et al*. 2016; Sperry *et al*. 2017), we propose a new instantaneous stomatal optimisation model that explicitly accounts for sapwood traits. During photosynthesis, CO_2_ is assimilated from the atmosphere through stomata in exchange for H_2_O, which is drawn from the soil to the leaves by negative pressures along water potential gradient. Negative water potentials impose costs on the plant through a heightened risk of xylem cavitation, subsequent loss of conductivity and cost of restoring conductivity (Choat *et al*. 2018). Given that plants must compromise between increasing photosynthesis and reducing hydraulic costs, the stomatal optimisation model seeks to to maximise net photosynthesis at any moment by balancing these opposing forces.

For a given height, trait, and environment, the stomatal model maximises the instantaneous net rate of photosynthesis for a given hydraulic cost, both of which vary as a function of the leaf water potential (*ψ*_leaf_):

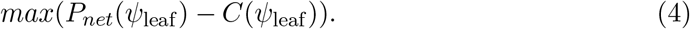

Here, *P*_*net*_ is the net photosynthetic gain after accounting for the leaf respiration rate and *C* is the hydraulic cost associated with the transpiration of water. *P*_*net*_ and *C* are functions of the leaf water potential (*ψ*_leaf_), which is directly related to the rate of stomatal conductance (see below). Eq. 4 is satisfied when

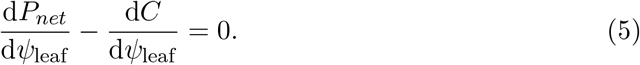

Stomatal conductance (*g*_*c*_) in this model is *a priori* unknown, and is an emergent outcome of the optimisation. *g*_*c*_ is directly linked to *ψ*_leaf_ under the assumption that instantaneous water supply, via xylem transport (*E*_supply_), must be equivalent to the atmospheric demand for water (*E*_demand_; Sperry & Love 2015; Sperry *et al*. 2017). Assuming no segmentation, *E*_supply_ depends on the water potential gradient between the soil and the leaf, given by the definite integral of the unique hydraulic vulnerability curve (Eq. 10), bounded by *ψ*_leaf_ and *ψ*_soil_:

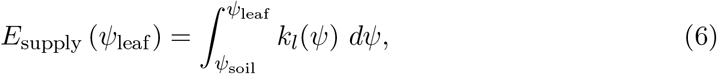

where leaf-specific hydraulic conductance (*k*_*l*_) varies with *ψ*. For a given *ψ*_soil_, *E*_supply_ increases with *ψ*_leaf_ but at a diminishing rate (Sperry *et al*. 2017).

Ignoring the leaf boundary layer conductance to water vapour, *E*_demand_ depends on the atmospheric vapour pressure deficit (*D*) and the rate of stomatal conductance of water vapour (*g*_*w*_ = 1.6 *g*_*c*_ ), which in turn varies with *ψ*_leaf_:

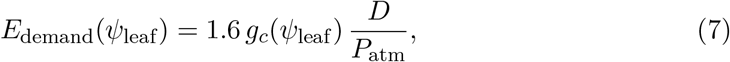

where 1.6 represents the molecular diffusion ratio of H_2_O to CO_2_. Setting *E*_demand_ = *E*_supply_ and rearranging shows how *g*_*c*_ varies as a function of *ψ*_leaf_ and other parameters:

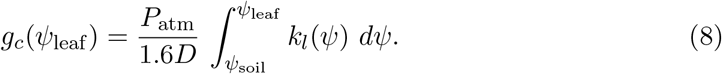

Importantly, Eq. 8 specifies a monotonic relationship between *g*_*c*_ and *ψ*_leaf_, meaning the optimisation in Eq. 5 could be performed with respect to either variable.

#### 3.2.1 Hydraulic costs

We implemented a novel representation of hydraulic costs (*C*), extending a previous cost function that expresses costs in rates of carbon loss per unit leaf area (Bartlett *et al*. 2019). In this model, *C* is assumed to account for the loss of conductivity in the xylem pathway incurred by negative *ψ*_leaf_. The hydraulic cost function therefore represents the absolute amount of carbon required to restore xylem conductivity.

Following Bartlett *et al*. (2019), we assume *C* increases with *ψ*_leaf_, as *k*_*l*_ declines from its maximum possible rate (*k*_l,max_):

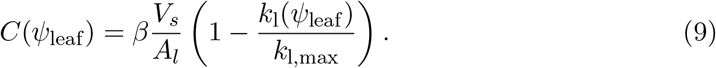

Here, *V*_*s*_ is the sapwood volume which is normalised by *A*_*l*_ to define *C* in units of leaf area and *β* is the carbon cost per unit of sapwood volume lost to embolism per unit time. Importantly, this formulation implies plants will experience carbon costs even when stomata are closed, which could act as a proxy for the effect of cuticular conductance of water from the leaf to the atmosphere (Choat *et al*. 2018).

*k*_*l*_ is assumed to decline from *k*_*l,max*_ as *ψ*_leaf_ becomes more negative following a Weibull distribution:

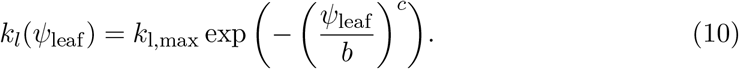

Here, *c*, and *b*, are the sensitivity and shape parameters of the Weibull distribution, respectively. Embolism damage is assumed to recover instantaneously as leaf water potential becomes less negative, with the carbon cost to repair this damage being accounted for in Eq. 3.

We made two extensions to Bartlett’s cost function (Eq. 9), to better capture carbon costs in relation to traits and size. First, sapwood volume per leaf area 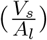 was calculated from our allometric model as:

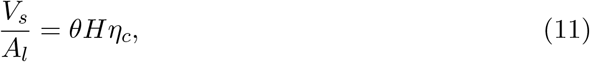

where *θ* is the sapwood to leaf area area ratio (also known as the Huber value) and *η*_*c*_ represents the average position of a leaf in the canopy as a proportion of the height of the plant (H).

Second, we developed a more mechanistic foundation for the rate parameter *β*. In Bartlett *et al*. (2019), *β* was parameterised based on an empirical correlation between the water potential of the transpiration pathway at which conductivity is halved (i.e. *P*_50_) and the amount of embolism that occurs at stomatal closure. There, *β* is assumed to increase with *P*_50_, representing the additional carbon investment in stem material required to increase drought tolerance. Instead, we assume that *β* is linked to wood density, *ρ*, based on the assumption that *ρ* is needed to translate a volume of wood into biomass, and that *ρ* additionally affects the risk of damage at a given pressure. This second assumption reflects empirical evidence showing that denser wood often confers greater resistance to embolism (Hacke *et al*. 2001; Janssen *et al*. 2020; Preston *et al*. 2006; Kiorapostolou *et al*. 2019; Hoffmann *et al*. 2011). These two effects are described by an equation with two parts

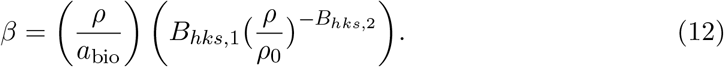

The first part of the right-hand side of Eq. 12 represents the fact that embolism would incur a greater loss of mass per unit sapwood volume. The second part of the right-hand side of Eq. 12 gives the rate of turnover, dependent on *ρ*, where *B*_*hks*,1_ is the average rate of sapwood turnover rate for a fully cavitated stem when *ρ* is equal to the global average, *ρ*_0_ (Table 2), and *B*_*hks*,2_ defines the strength of the trade-off between *ρ* and *β*. Thus, although *β* is assumed to increase linearly with *ρ* owing to the first term on the right-hand side of Eq. 12, *β* is also assumed to decline exponentially with increasing *ρ*, representing a greater resistance to embolism-inducing water potentials (Hacke *et al*. 2001).

Finally, we parameterise *k*_*l*,max_ by normalising the maximum sapwood-specific conductivity, *K*_*s,max*_ by *θ* and dividing by the path length of the conducting tissue (Xu *et al*. 2021):

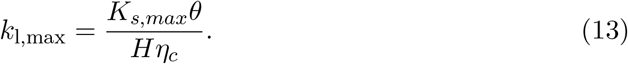

#### 3.2.2 Photosynthesis

Photosynthesis (*P*_*net*_) was modelled using a standard coupled stomatal–photosynthesis model, based on Fick’s first law of diffusion:

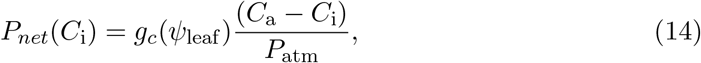

where *C*_a_ is the atmospheric concentration of CO_2_, *C*_i_ is the intercellular concentration of CO_2_, P_atm_ is the atmospheric pressure, and *g*_*c*_ is the rate of stomatal conductance to CO_2_. *C*_i_ is *a priori* unknown, but can be found by combining Eq. 14 with the Farquhar-von Caemmerer-Berry biochemical photosynthesis model (see Supporting Information) and numerically solving for *C*_i_.

### 3.3 Trait-based trade-offs

In order for trait-environment gradients to emerge, traits must capture trade-offs in ecological function with costs and benefits that vary with respect to the environment. These trade-offs are discussed below.

#### LMA, *ϕ*

Increasing *ϕ* reduces the rate of leaf area deployment for a given unit of biomass growth via its effect on the construction cost of leaves 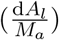 but also reduces the rate of leaf turnover (*t*_*l*_) according to the well-recognised leaf economics spectrum (Wright *et al*. 2004):

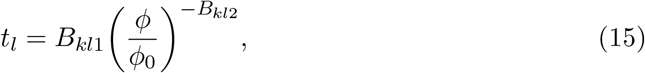

where *ϕ*_0_ is the global average of *ϕ* (see Table 2), *B*_*kl*1_ is the leaf turnover rate at *ϕ*_0_ and *B*_*kl*2_ is the steepness of the relationship between LMA and the turnover rate on a log-log scale. For *ϕ*, and other traits, the global average value is included such that variation in the magnitude of the trade-off exponent causes the trade-off axis to rotate upon a central trait value.

#### Maximum sapwood-specific conductivity, *K*_*s,max*_

*k*_l,max_ increases with *K*_*s,max*_ which increases the rate of water transport. However, sapwood with a higher *K*_*s,max*_ is also assumed to be less resistant to embolism, consistent with the hypothesis of a hydraulic safety-efficiency trade-off (Liu *et al*. 2019, 2021; Franklin *et al*. 2023):

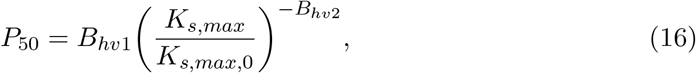

where *P*_50_ is the xylem pressure at which 50% of conductivity is lost, *B*_*hv*1_ is the *P*_50_ at the global average of *K*_*s,max*_ (*K*_*s,max*,0_; Table 2) and *B*_*hv*2_ is the steepness of the relationship between *K*_*s,max*_ and *P*_50_ on a log-log scale. *B*_*hv*1_ and *B*_*hv*2_ were parameterised based on the data presented in Liu *et al*. (2019).

*P*_50_, in turn, determines the sensitivity parameter of the hydraulic vulnerability curve, *b*:

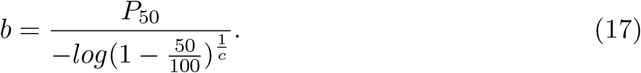

The shape parameter (*c*) might also vary with *K*_*s,max*_ and *P*_50_, perhaps to capture a trade-off between the width of the hydraulic safety margin (e.g. P50 - P88) and the maintenance of high conductivity under low soil moisture stress. However, as the validity of such covariation remains unclear, we instead set it as a constant value (Table 2).

#### Huber value, *θ*

The cross-sectional sapwood area to leaf area ratio (*θ*), influences growth rates in a number of ways. *k*_l,max_ increases with *θ* by increasing the cross-sectional area of sapwood supplying a given area of leaf with water. However, 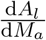 declines with *θ* due to its effect on the construction cost of sapwood and bark (see Supplementary Information). In addition, *C* increases with *θ* via its effect on *V*_*s*_ because a given unit of xylem damage incurs a greater absolute loss of carbon in plants with a greater amount of sapwood in our model.

#### Wood density, *ρ*

Increasing *ρ* reduces 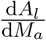 via its effect on the construction cost of sapwood and bark but also reduces the rate of sapwood damage per unit of conductivity loss as described in Eq. 12.

### 3.4 Implementation and trait optimisation

Plant growth was simulated in the **plant** R package (Falster *et al*. 2016) using the FF16w physiological module. The FF16w physiological module adds the stomatal sub-model described above into the base FF16 module. Height growth rates were numerically solved across gradients of soil water potential (*ψ*_soil_), vapour pressure deficit (*D*), atmospheric CO_2_ concentration (*C*_*a*_) and above-canopy photon flux density (*I*_0_) as well as at different plant heights. When simulating across environmental gradients, height growth rates were calculated for 1m tall plants. To find the value of a traits optimising height-growth rates in a given environment, we minimised the objective function: 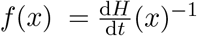. When considering a single trait, we used the **optimise** function from the **stats** R package. When considering multiple traits, we used generalised simulated annealing, as implemented in the **GenSA** R package (Xiang *et al*. 2013), using a maximum of 5000 iterations of the search algorithm.

#### 3.4.1 Finding the optimum *ψ*_leaf_

At each time step, we solved for the optimum *ψ*_leaf_ using numerical methods, as no analytical solution for the *ψ*_leaf_ that satisfies Eq. 5 exists. We solved Eq. 5 using the golden-section search algorithm which requires upper and lower boundary values (Kiefer 1953). For the upper boundary value, we assumed that *ψ*_leaf_ cannot be less negative than *ψ*_soil_. For the lower boundary value, we assumed that *ψ*_leaf_ could not be more negative than the critical point at which 99% of hydraulic conductivity is lost (*ψ*_crit_).

### 3.5 Parameterisation

In addition to any sources described above, extra parameters in the FF16w module were either obtained from existing literature or were parameterised where data was available to represent a self-supporting woody plant using the AusTraits trait database (Table 2), which is a collation of Australian plant trait data and is the single largest collation of trait data within a single continent (Falster *et al*. 2021). We considered our use of the AusTraits database appropriate given our focus in the present study on simulating qualitative predictions of trait-environment relationships, rather than matching site or species-specific observations. Along these lines, for parameters which relatively little is known, such as B_hks,1_, we simply selected values which yielded realistic simulations of trait optima. Otherwise, the model was parameterised largely using the default parameters available in the base FF16 module of **plant** (Falster *et al*. 2016).

## 4 Results

### 4.1 Emergent response of stomatal behaviour to traits and environment

Under our optimality framework, sensitivity of stomatal conductance to the environment is an emergent property rather than the outcome of an empirical relationship.

In line with empirical observations, our stomatal optimisation model predicts a depression in *g*_*s*_ (Figure 1) shortly after midday due to a peak in the atmospheric vapour pressure deficit (Koyama & Takemoto 2014), as well as a downregulation of *g*_*s*_ as soils dry (Zhou *et al*. 2013). Considering univariate responses of *g*_*s*_ to the environment, our model predicts a decline in *g*_*s*_ with increasing soil water potential (*ψ*_soil_), vapour pressure deficit (*D*) and atmospheric CO_2_ concentration (*C*_*a*_), but an increase with above-canopy photon flux density (*I*_0_). Moreover, traits influence the response of *g*_*s*_ to the environment; plants with greater *K*_*s,max*_ and *θ* achieve a higher *g*_*s*_ in a given environment, and are more water profligate (Figure 1).

**Figure 1:**
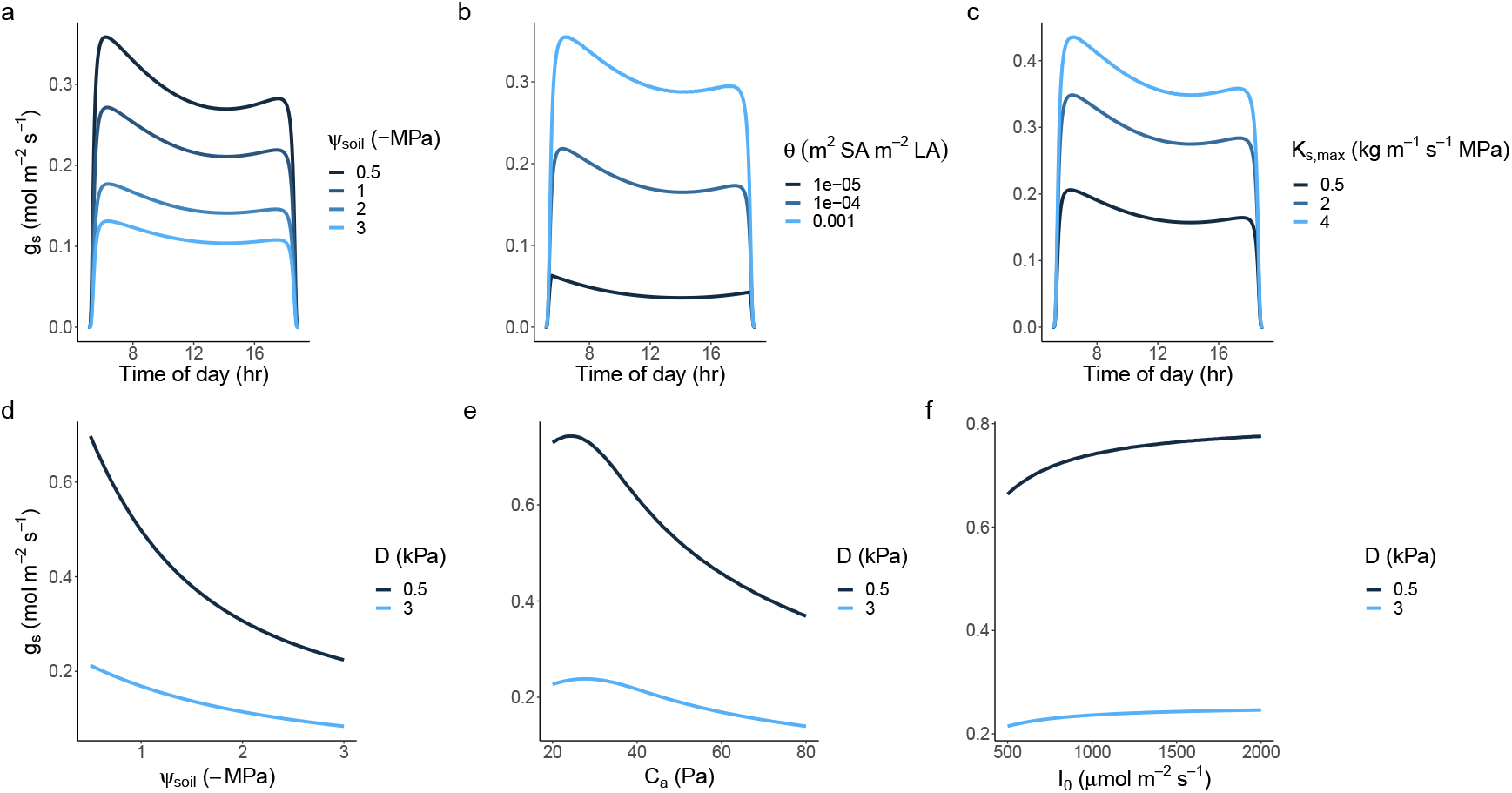
The sensitivity of stomatal conductance, *g*_*s*_, to the environment and traits emerges from our stomatal optimisation framework. The model successfully predicts stomatal response to diurnal fluctuation in atmospheric vapour pressure deficit (D) and light availability (I_0_) as **a)** soils dry, with **b)** increasing sapwood conductivity, *K*_*s,max*_ and **c)** sapwood to leaf area allocation, *θ*. The model also predicts that stomatal closure occurs as **(d-f)** *ψ*_soil_, *C*_a_ and I_0_ decreases and as the air becomes drier.

### 4.2 Emergent response of height growth rate to the environment

For a given set of traits at a fixed height, absolute height growth rates were greater in wetter soils (i.e. less negative *ψ*_soil_), lower *D*, higher *C*_*a*_ and higher *I*_0_ (as represented by the contour lines in Figure 2).

**Figure 2:**
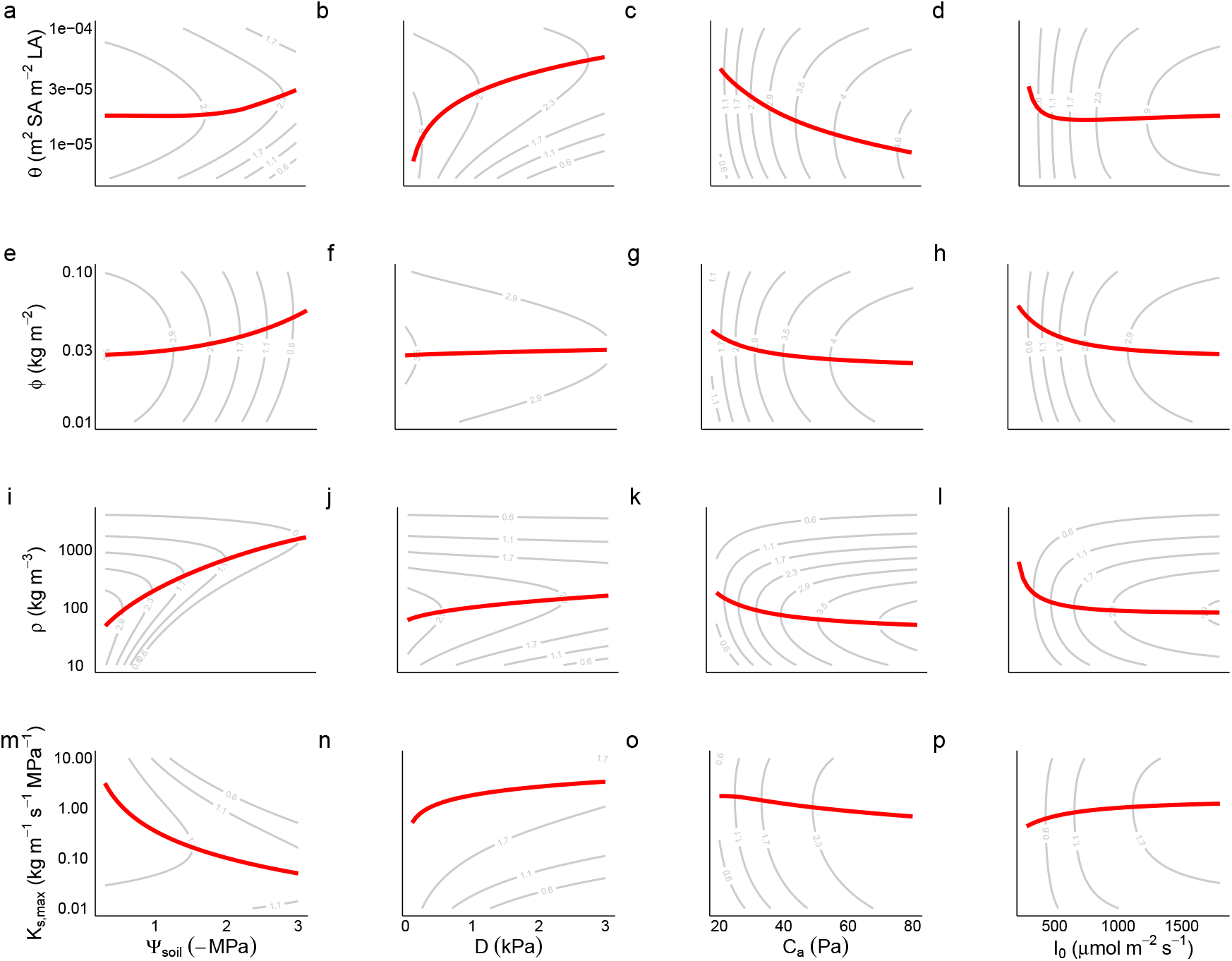
Predicted environmental sensitivity of four traits. Red lines connect the trait values optimising height-growth rate across fifty simulated points for each environmental gradient. Traits were optimised one at a time with all other traits held at their default value. Environmental variables were also varied one at a time, with all other variables being held at constant values: *ψ*_soil_ = 0.5 -MPa, D = 0.5 kPa, C_*a*_ = 40 Pa, and I_0_ = 1800 *χ*mol m^*−*2^ s^*−*1^. The grey countours represent height growth rates for a 1m tall plant.

### 4.3 Individual trait responses to the environment

#### Optimal Huber value, *θ*

Increasing *θ* entails an increase in the water transport rate per leaf area at the cost of a reduced efficiency of leaf area deployment. Moreover, greater *θ* leads to greater volume-based hydraulic costs and mass-based respiration and turnover rates. Thus, in general, *θ* is expected to become larger where the benefit of a greater *k*_*l,max*_ to 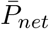 are more apparent relative to the combined costs of construction, respiration and tissue turnover. Specifically, we found that the response of *θ* to *ψ*_soil_ was non-monotonic (leading to the flat response in wetter soils in Figure 2a), especially at greater heights (e.g. by following the horizontal gradient towards the top of Figure 3a), and this was because the 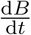 optimising *θ* initially declined as soils dried, before increasing again in very dry soil (Figure 4a). The initial decline occurred because it was more profitable for plants to obtain water for transpiration by making *ψ*_leaf_ more negative and saving on instantaneous costs associated with greater sapwood volume. In very dry soils, however, *θ* increased again because *ψ*_*soil*_ approached the critical xylem pressure, *ψ*_crit_, such that it was more profitable to supply the transpiration stream by increasing sapwood volume (Figure S5). *θ* also increased with *D* and declined with *C*_*a*_ (Figure 2b,c). In the former case, this increase in the atmospheric demand for water induced by *D* (Eq. 7) was compensated for by an increase in the water transport rate via increasing *θ*. In the latter case, increasing C_a_ caused partial stomatal closure (i.e. lower *g*_*s*_; Figure 1) and a reduction in *E*_demand_ which was achieved by reducing *θ*.

**Figure 3:**
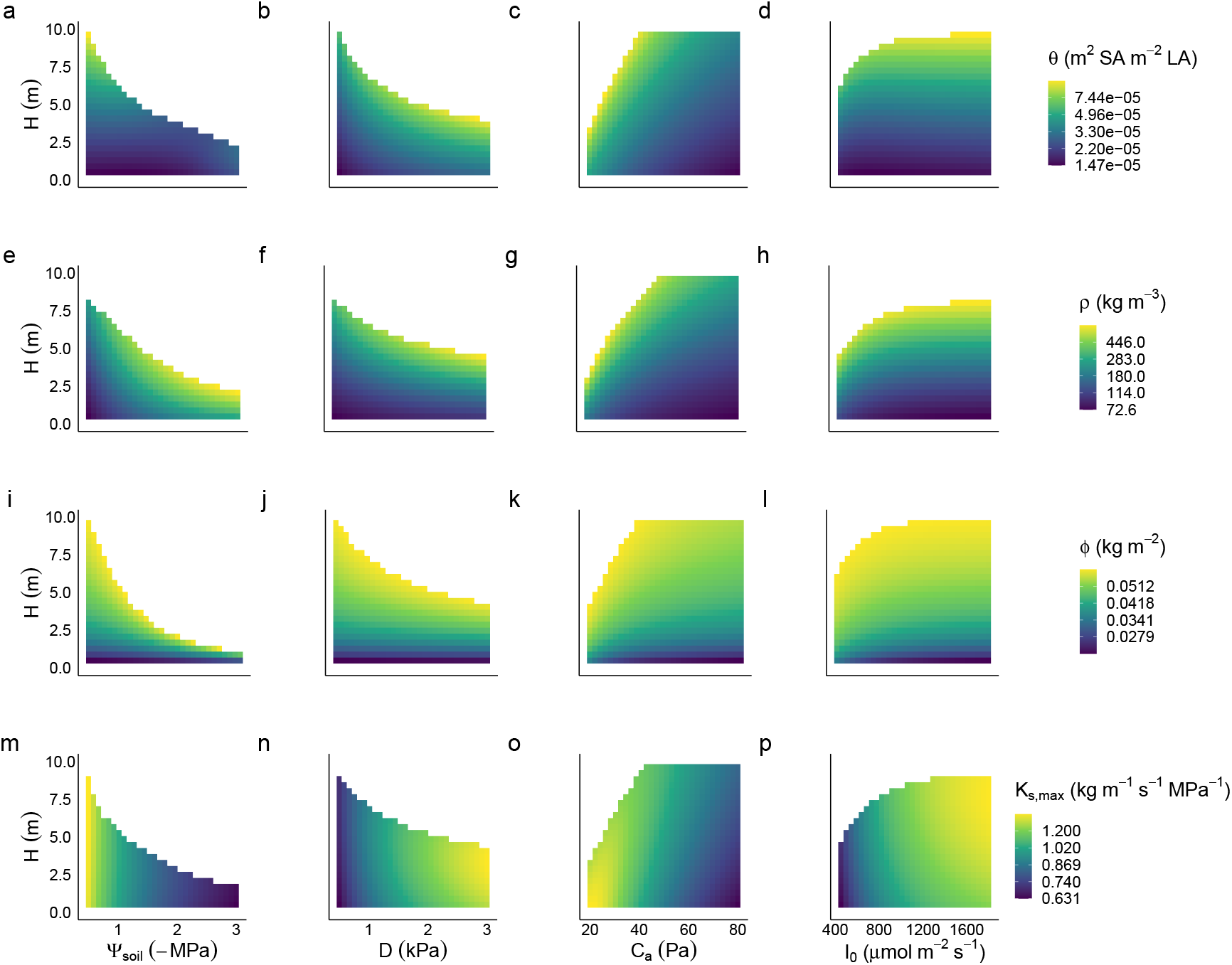
Predicted ontogenetic shifts in traits. Cell shading represents the value of the height-growth optimising trait for a plant of a given height and under a given set of environmental conditions. Traits were optimised one at a time with all other traits held at their default value. Environmental variables were also varied one at a time, with all other variables being held at constant values: *ψ*_soil_ = 0.5 -MPa, D = 0.5 kPa, C_*a*_ = 40 Pa, and I_0_ = 1800 *χ*mol m^*−*2^ s^*−*1^. White space in each panel indicates positions in the environment-height space under which no trait conferred positive height growth rates.

**Figure 4:**
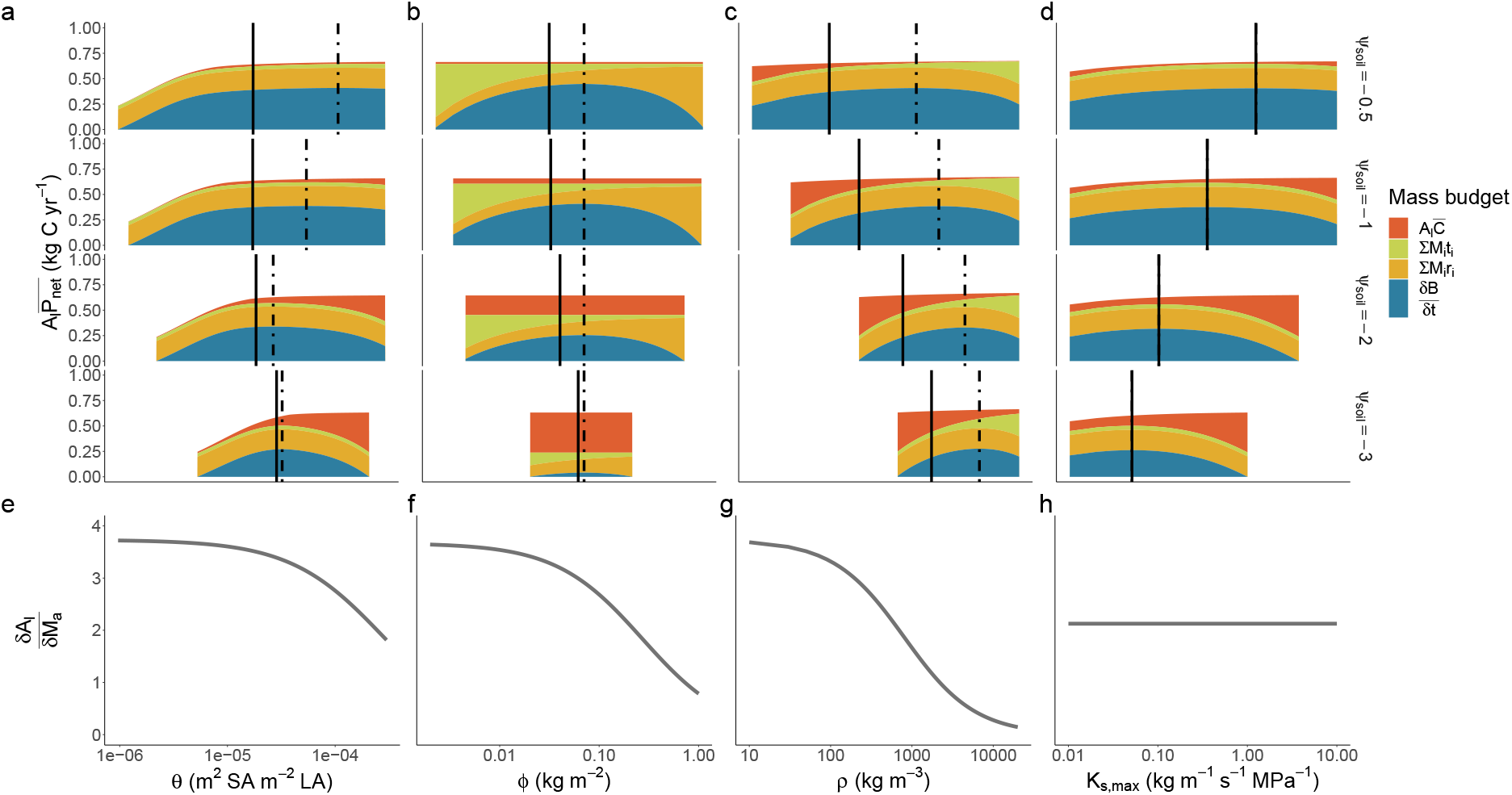
Decomposition of the components determining trait optima across a soil water availability (*ψ*_*soil*_) gradient. The top row of panels shows how net biomass production,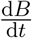 emerges for each trait across a declining soil water availability gradient (i.e. from the top to the bottom of panels **a-d**) as the residual of total assimilation, 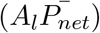, after accounting for hydraulic The costs, 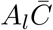, turnoverΣ*M*_*i*_*t*_*i*_ and respiration Σ*M*_*i*_*r*_*i*_ of each plant tissue, *i*. The trait value maximising 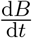 is indicated by the dashed vertical bar. The solid vertical line indicates the trait value maximising the height-growth rate, 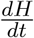. The 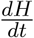.optima emerges through multiplication of 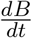 with the rate of leaf area deployment per unit of live mass growth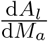, shown in the second column. For most traits, this causes the 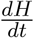 optima to be lower than the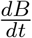 optima, owing to the greater value of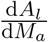at low trait values in panels **e-g** but also explains why the optima are equivalent for *K*_*s,max*_.

The non-monotonic response of the height-growth optimising *θ* to *I*_0_ in Figure 2 emerged despite the fact that the *θ* optimising 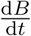 responded monotonically positive to *I*_0_ (Figure S2a). This was because, under very low light, 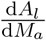 causes the height-growth optima to shift more strongly towards the value of *θ* optimising 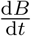 (Figure S2a,e) than it does under high light conditions.

#### Optimal LMA, *ϕ*

*ϕ* increased with environmental harshness, namely increasing with soil and atmospheric aridity but declining with increasing atmospheric CO_2_ and light availability (Figure 2).

In our model, leaf respiration increases with *ϕ* while leaf turnover decreases, yielding a hump-shaped relationship between *ϕ* and 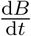. However, because *ϕ* does not mediate the response of 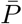 to the environment (Figure 4), the *ϕ* maximising 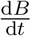 is also invariant to the environment. Nevertheless, *ϕ*-environment relationships emerge as changes in the absolute value of 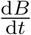n and 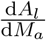, which decreases monotonically with *ϕ*. Put simply, the effect of low *ϕ* on construction outweighs the combined carbon losses to respiration and turnover when 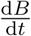 is high.

#### Optimal Wood density, *ρ*

Increasing *ρ* reduces the maximum rate of embolism incurred by a given proportional loss of conductivity (Eq. 12) but also incurs an increasing construction cost of sapwood and bark. Thus, *ρ* was predicted to be larger in more arid environments or environments with lower atmospheric CO_2_ (Figure 2) where more negative optimal *ψ*_leaf_, and thus embolism risk, are encountered. Counter-intuitively, even though the optimal *ψ*_leaf_ also becomes more negative as *g*_*s*_ and *A*_*net*_ increase with light availability, we found that *ρ* declines with light availability. This occurred because, similarly to *ϕ* and *θ*, a reduction in 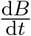 under low light caused the 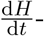 optimising *ρ* to shift more strongly towards the 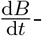 optimising *ρ* (Figure S2c,g).

#### Optimal Maximum sapwood-specific conductivity

*K*_*s,max*_: *K*_*s,max*_ directly tradedoff with *P*_50_ such that, all else being equal, a plant with a higher *K*_*s,max*_ had a greater *k*_*l,max*_ but experienced greater embolism at less negative *ψ*_leaf_. Importantly, because *K*_*s,max*_ does not influence 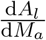, the *K*_*s,max*_ optimising 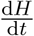 is equivalent to the *K*_*s,max*_ optimising 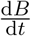 (Figure 4). We observed contrasting responses of *K*_*s,max*_ to *ψ*_soil_ and *D*, with *K*_*s,max*_ declining in the former case and increasing in the latter case (Figure 2). Contrasting responses emerged because variation in *K*_*s,max*_ impacted the net carbon assimilation and hydraulic cost differently. In the case of *ψ*_soil_, the predicted response emerged because hydraulic cost rose more rapidly as *K*_*s,max*_ increased in drier soils, such that maximum leaf-level 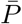 occurred at lower *K*_*s,max*_ (Figure S4). In other words, *K*_*s,max*_ was predicted to decline to improve embolism resistance in the xylem in moisture-stressed locations. In the case of *D*, the predicted increasing response emerged because the marginal benefit to *A*_*net*_ of increasing *K*_*s,max*_ was much greater under high atmospheric aridity than low, meaning that maximum leaf-level 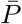 occurred at higher *K*_*s,max*_ (Figure S4). In other words, for plants experiencing dry air, more rapid water transport is more beneficial to growth, provided sufficient soil water is available to maintain the transpiration stream.

### 4.4 Individual trait responses to height

In our model, individuals do not experience competition and thus increasing height has no effect on the environmental conditions experienced by plants. Instead, the response of traits to ontogeny primarily emerges through the effect of height on 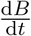 on a per-leaf basis (i.e. the total biomass growth of the plant divided by *A*_*l*_). As height increases, the cost of respiration and turnover in non-photosynthetic tissues as a fraction of leaf-level assimilation also increases. Moreover, the hydraulic cost of water transport increases with the sapwood volume as height increases, all else being equal. Thus, in our model, increasing height can be viewed analogously as a shift towards an increasingly unproductive environment.

It follows from our above analysis of trait-environment predictions, then, that for traits influencing the relative allocation and construction cost of plant tissues (i.e. *θ, ρ, ϕ*), increasing height caused the height-growth optimising trait value to move towards the construction of more expensive tissues (e.g. more dense wood; Figure 3a-l, viewing each subpanel vertically) which minimise losses to tissue turnover.

The very minor, yet positive, response of *K*_*s,max*_ to height has a more simple explanation (Figure 3m-p). In our model, *K*_*s,max*_ does not influence the construction cost of tissues and therefore exerts influence on height growth rates exclusively through 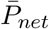. As plants grow taller, *k*_*l,max*_ declines, reflecting an increase in the length of the hydraulic pathway (Eq. 13). Thus, plants are predicted to compensate for greater heights by increasing *K*_*s,max*_. Opposing this shift towards greater *K*_*s,max*_ is the fact that taller plants experience greater damage to the hydraulic pathway per unit of lost conductivity (Eq. 9), and must therefore balance the benefit of more rapid water transport against the need for safer xylem (i.e. a more negative P_50_) at greater heights, explaining the low sensitivity of this trait to plant height.

### 4.5 Species turnover across soil moisture gradients

In the analyses above, we have explored how trait optima shift across environmental gradients considering one trait at a time. However, selection operates on whole-plant fitness integrated across multiple traits. Using the same framework as above, we jointly optimised height-growth rate across the four traits. In general, traits responded in the same direction as when optimised in isolation, but we observed much greater sensitivity in *K*_*s,max*_ and a much reduced sensitivity in the remaining traits (Figure 5).

**Figure 5:**
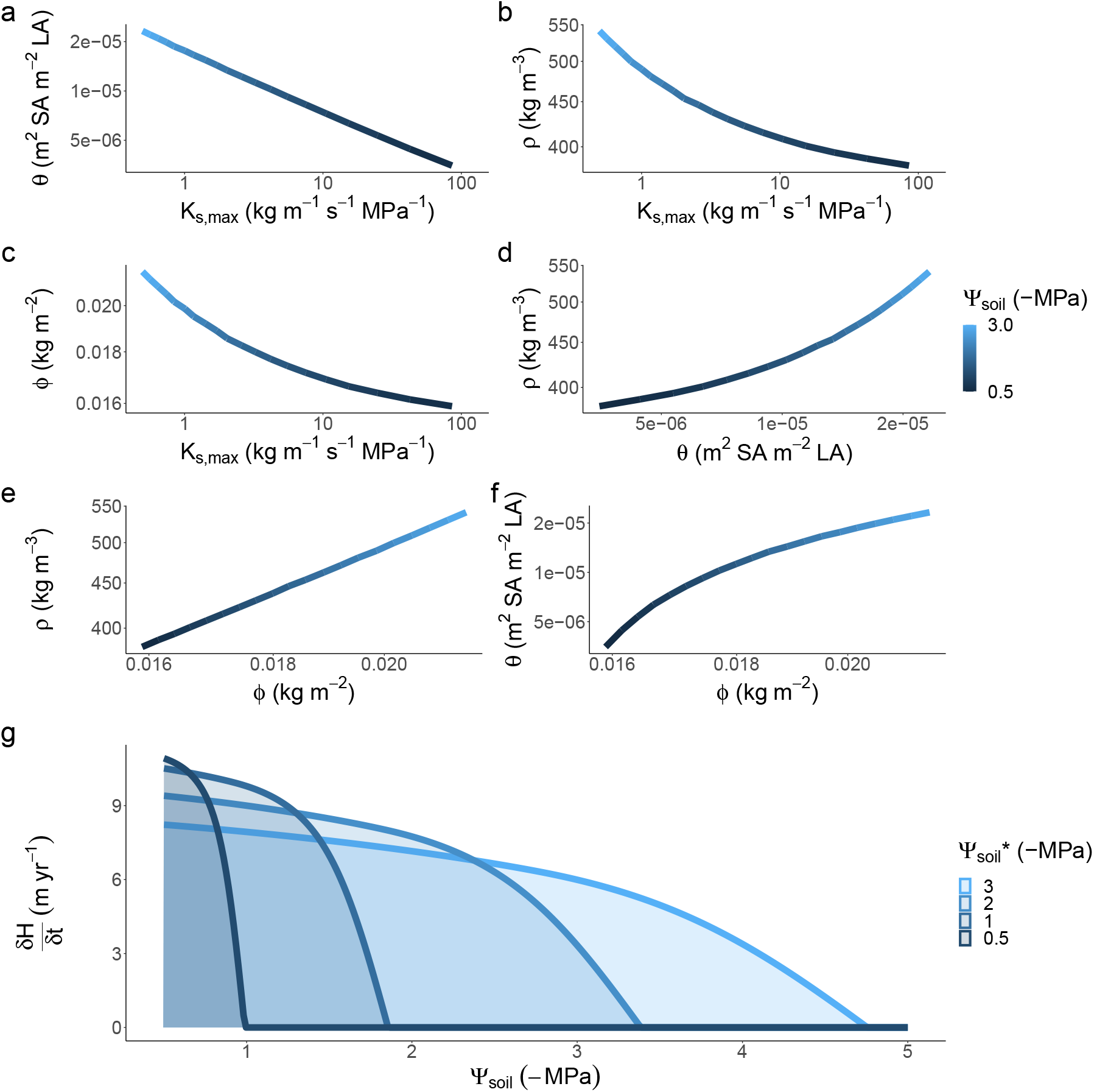
Species turnover across a soil moisture gradient emerges from height-growth rate optimisation of multiple traits simultaneously. Joint optimisation of four traits, being the sapwood to leaf area ratio (*θ*), wood density (*ρ*), leaf mass per area (*ϕ*) and sapwood conductivity (*K*_*s,max*_) reveals a coordinated shift towards drought-resilient traits in drier environment (i.e. higher *θ, ϕ* and *ρ* and lower *K*_*s,max*_). Generating height-growth rate curves for four hypothetical species optimised to different points along the soil moisture gradient (i.e. 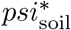 reveals a trade-off between the maximum achievable growth rate and tolerance for dry soils).

Our optimisation approach can also yield insights into how the emergent properties of plant communities respond to turnover in plant strategies across soil water availability gradients. We found that, while all plants grow faster when water is more plentiful (Figure 5), plants that are optimised to grow in wet soils achieve the highest growth rates. This is because they have traits which maximise water transport rates and minimise tissues construction costs, including high *K*_*s,max*_ and low *ϕ, ρ* and *θ*. However, species with such traits are also predicted to have narrow fundamental niches (as defined by the range of conditions where they can maintain positive growth), because their growth rate declines rapidly as soils dry. In drier soils, species with low *K*_*s,max*_ and high *ϕ, ρ* and *θ* are expected to dominate because they can maintain positive growth rates even under drought (i.e. have wider environmental tolerances; Figure 5).

## 5 Discussion

Recent advances in plant optimality theory that link stomatal behaviour and gas exchange with plant hydraulics are enabling a mechanistic link between plant traits and the environment. By integrating a stomatal optimisation criterion into an existing height-growth model, we made predictions for how four key traits (LMA, SA:LA, sapwood-specific conductivity and wood density) should shift to optimise plant performance over a range of environmental gradients, including soil moisture. Broadly speaking, these predictions qualitatively match empirical trait-environment patterns (Table 1). Moreover, we predicted how these traits should shift throughout individual ontogeny. Our framework establishes the groundwork for future modelling work seeking to understand how the trait composition of vegetation, and thus, ecosystem processes, are likely to respond to future changes in water availability (Sakschewski *et al*. 2015; Harrison *et al*. 2021; Feeley & Zuleta 2022).

### 5.1 Trait responses to soil moisture

We predicted an increase in wood density as soil moisture declined in line with many (Onoda *et al*. 2010; Pickup *et al*. 2005; Towers *et al*. 2023; Sakschewski *et al*. 2015), but not all (Swenson & Enquist 2007; Chave *et al*. 2009) large scale empirical studies. This prediction emerges from our modification of the hydraulic cost function implemented in Bartlett *et al*. (2019), whereby we assumed that permanent damage to the xylem imposed by xylem cavitation, *β* declines with wood density, which is consistent with empirical evidence showing that denser wood has a higher resistance to xylem implosion (Hacke *et al*. 2001). In principle, the same outcome would emerge if wood density instead had a negative relationship with *P*_50_; the critical process underlying the predicted response of wood density to soil moisture is that minimising hydraulic costs becomes more beneficial relative to minimising stem construction costs in drier conditions.

As soils dried, wood density for 1m tall seedlings was predicted to increase 35-fold from 47.8g cm^*−*3^ to 1651g cm^*−*3^, closely matching the smallest and largest observed values in nature but far exceeding observed fold-changes at a community level (Figure 6).

**Figure 6:**
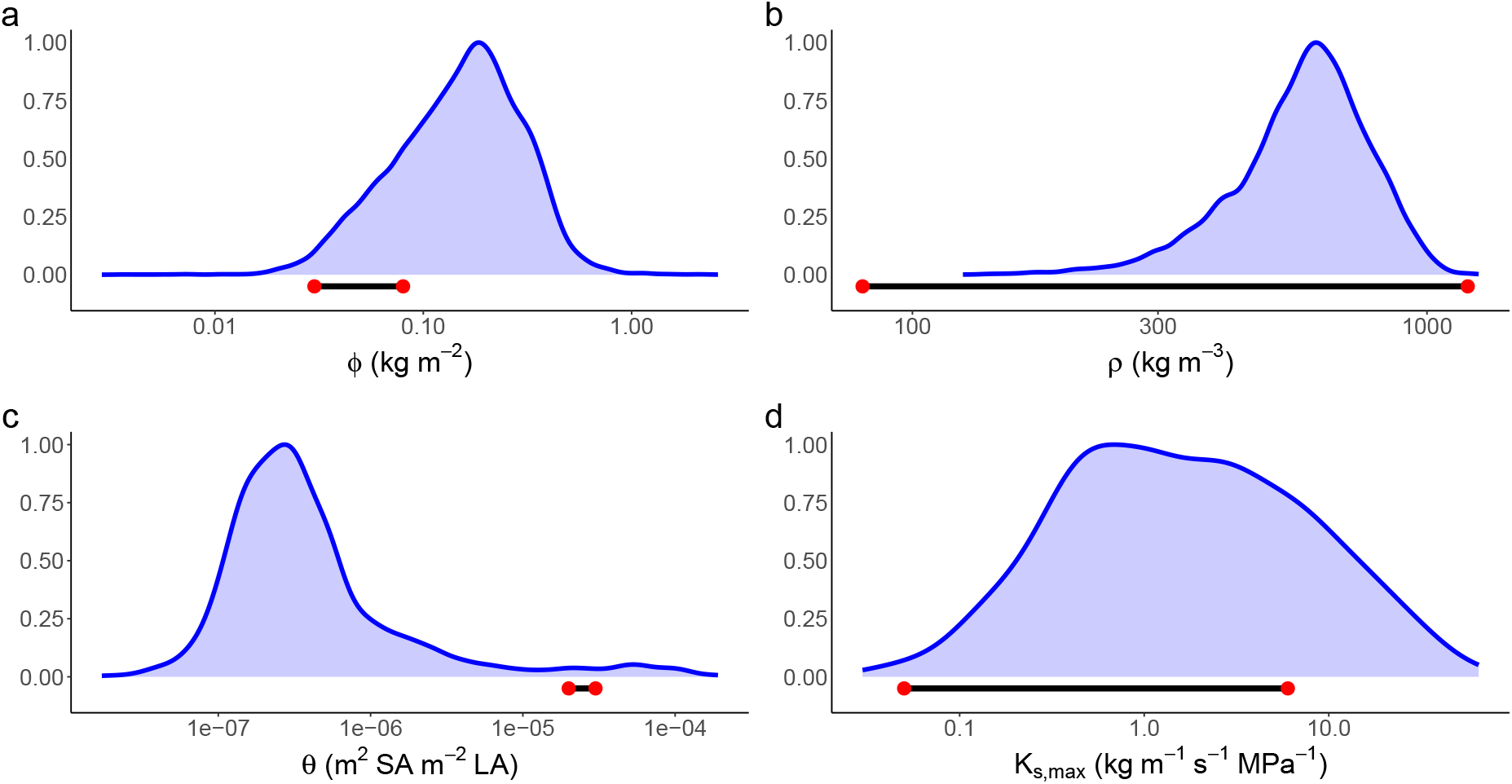
Simulated focal trait sensitivity to soil moisture relative to the distribution of empirical observations. The red points represent the minimum and maximum simulated value of each trait across the soil moisture gradient depicted in Figure 2. Blue fields are the scaled density plots of empirical observations for each of the four focal traits for woody plants in the AusTraits trait database (Falster *et al*. 2021).

Our primary goal in this paper was to explore directional relationships in traits based on simplified trait trade-offs, and in this sense, we have captured empirical patterns. However, the exaggerated response of *ρ* to environment suggests key processes that are currently missing in the model may help constrain the range of predicted values. For example, the risk of stem breakage due to wind load would put upward selective pressure on wood density (Rifai *et al*. 2016). Similarly, if denser wood is more resistant to stem damage inflicted by pathogen attack and other causes, this would also cause wood density to be larger (Larjavaara & Muller-Landau (2010) and references within). The high values of wood density that we observed, especially for taller plants, probably relate to uncertainty regarding the parameters *B*_*hks*,1_ and *−B*_*hks*,2_, which represent the cavitation-related maximum yearly rate of sapwood loss and the trade-off slope between *B*_*hks*,1_ and wood density, respectively. Indeed, although much is known about how xylem conductance relates to stem water potential, little is known about how loss of conductivity translates into a holistic cost to the plant in absolute carbon units (Bartlett *et al*. 2019; Potkay & Feng 2023), and in this case we simply selected a value of *B*_*hks*,1_ which best captured realistic values of the four focal traits (Figure 6).

In a similar manner to wood density, our model predicted a shift towards denser, more expensive leaves (i.e. higher LMA) in drier soils. This outcome is consistent with empirical studies at a variety of spatial scales (Dwyer *et al*. 2014; Wright *et al*. 2004; Towers *et al*. 2023; Niinemets 2001). Notably, the predicted response of LMA to soil moisture emerges without influencing the plant hydraulic pathway. Instead, it emerged from the well-recognised trade-off between leaf dry mass per area and the rate of leaf turnover (Wright *et al*. 2004). Put simply, when photosynthetic rates decline across a drying soil moisture gradient, plants are incentivised to invest carbon in longer-lasting tissues to avoid frequent leaf turnover, even if this leads to a more expensive up-front cost to growth. In reality, LMA could also influence plant hydraulics in addition to the above-mentioned effect and this could help to explain the relatively low sensitivity that we observed in this trait (Figure 6). For example, while we observed an approximately two-fold increase in LMA for a 1m tall plant along the simulated soil moisture gradient, a 10-fold shift in mean LMA has been observed across a continental rainfall gradient in Australia (Towers *et al*. 2023)). Possible mechanisms include the involvement of LMA in maintaining leaf rigidity under dry conditions (Poorter *et al*. 2009) as well as determining the conductivity of the leaf hydraulic pathway (Simonin *et al*. 2012). Additionally, higher light in harsh environments selects for higher leaf nitrogen per area, and thus LMA (Dong *et al*. 2017), an effect not captured in our current formulation.

Maximum sapwood-specific conductivity declined with a drying soil, as expected from theory and directly encoded in a trade-off between sapwood conductivity (i.e. sapwood efficiency) and vulnerability of the xylem to cavitation (Eq. 16). Evidence for the hydraulic safety-efficiency trade-off is mixed, with some studies demonstrating moderate correlations between maximum sapwood-specific conductivity and *P*_50_ (Liu *et al*. 2019, 2021) whereas others demonstrate little to no correlation between these variables (Gleason *et al*. 2016). However, triangular distributions in empirical data demonstrates that, while species can exhibit a range of combinations of maximum sapwood-specific conductivity and *P*_50_, achieving both high safety and efficiency appears very uncommon, implying the presence an upper boundary to the combined value of these traits (Gleason *et al*. 2016; Liu *et al*. 2021). As for species which fall towards the lower corner of the safety-efficiency space (i.e. low safety and efficiency), emerging evidence suggests that these taxa may occur more often in mesic, non-seasonal environments, because neither high maximum sapwood-specific conductivity nor *P*_50_ are required in these locations (Liu *et al*. 2021). Thus, our prediction may be more applicable for seasonal climates where selection co-optimises these traits.

Another important consideration is that while most analyses of the hydraulic safety-efficiency trade-off, including this analysis, focus on xylem efficiency at the sapwood-level, recent evidence suggests that the trade-off is mediated instead at the individual conduit level (Franklin *et al*. 2023). The implication of this is that the safety-efficiency trade-off may be weaker at the sapwood-level because maximum sapwood-specific conductivity is determined by both the conductivity of an individual conduit (*K*_*c*_) and the number of conduits, which can vary independently (i.e. by moving along both the S and F axis described in Zanne *et al*. 2010). Conceptually, then, our model can be considered a special case of optimising *K*_*c*_ when the lumen fraction is fixed. Further modelling work could investigate a link between maximum sapwood-specific conductivity and plant constructions costs in addition to the vulnerability of xylem to cavitation.

Our analysis of SA:LA builds on existing work investigating the theoretical relationship between this trait and *ψ*_soil_ (Westoby *et al*. 2012). Westoby *et al*. (2012) predicted that SA:LA responds in a hump-shaped manner to soil water availability because the initial benefit to plant revenue of increasing sapwood area to maintain the transpiration stream is countered by the increasing resistivity of soil as *ψ*_soil_ continues to decline. Our study also reveals a non-monotonic response of SA:LA to *ψ*_soil_ but for different reasons. Against our expectation, the SA:LA maximising 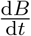 initially declined as soils dried because plants adapted by reducing sapwood area and making *ψ*_leaf_ more negative, enlarging the sapwood area only when *ψ*_soil_ approached *ψ*_crit_ at the driest end of the gradient. The reason why this occurred is because our model assumed that hydraulic costs are proportional to sapwood volume. In other words, a loss of conductivity in a larger sapwood volume is assumed to represent embolism in a greater absolute number of conduits. As such, the potential marginal benefit of a greater sapwood area in drier conditions was more than offset by greater hydraulic costs. Multiplication of proportional conductivity loss by sapwood volume was a chosen aspect of our representation of the hydraulic cost function in carbon units, but this implicitly assumes that hydraulic costs are associated with rebuilding the xylem (Gauthey *et al*. 2022). Hydraulic recovery may, however, be achieved via other mechanisms including bubble dissolution and xylem refilling (Klein *et al*. 2018) which, if less costly than rebuilding xylem, would potentially increase the selective advantage of higher SA:LA in drier environments, as is observed in nature (Towers *et al*. 2023), although we note that evidence for these mechanisms remains limited.

### 5.2 Trait responses to other environmental variables

A key outcome of our model was that it predicted the diverging response of maximum sapwood-specific conductivity to increasing soil dryness (i.e. declining) and atmospheric aridity (i.e. increasing) that is often observed in nature (Olson *et al*. 2020; Gleason *et al*. 2013). Increasing maximum sapwood-specific conductivity with atmospheric aridity emerged as a prediction from our model because it was more economical for plants to satisfy the additional evaporative demand for water by increasing the sapflow rate and minimising the water potential gradient between soil and the canopy. For brevity we do not consider the interactive effect of environmental variables in this analysis. However, as hypothesised in other studies (e.g. Gleason *et al*. 2013), the positive response of maximum sapwood-specific conductivity to atmospheric aridity is likely to be strongest when soil water is plentiful and the cost of low embolism resistance (due to a less negative P_50_) is minimal. It is less clear, how maximum sapwood-specific conductivity would respond to the combined effect of increasing soil and atmospheric dryness where the importance of embolism resistance conferred by more negative P_50_ becomes more prominent. Nevertheless, given that these conditions are likely to emerge in some locations under climate change, it is is worthy of further investigation.

Although our model adequately predicted the qualitative response of LMA to soil moisture observed in nature, it yielded contrasting responses to nature for light availability (Poorter *et al*. 2009; Ellsworth & Reich 1992; Neyret *et al*. 2016). There are number of possible reasons for this discrepancy. Firstly, it has been suggested that declining LMA towards lower light conditions reflects a shifting allocation of leaf biomass towards a greater deployment of leaf area so as to maximise light interception (Poorter *et al*. 2009). In our model, however, total leaf area is fixed to plant height and, as such, there is no benefit to light interception conferred to the plant by declining LMA. Secondly, and perhaps more importantly, we did not consider adaptive responses in leaf photosynthetic capacity which would cause the optimum LMA to increase with light availability as plants invest in greater photosynthetic capacity per unit area to better utilise available light (Poorter *et al*. 2009; Dong *et al*. 2022).

### 5.3 Trait responses to height

Our model accurately predicted the qualitative direction of observed trait responses to ontogenetic shifts in plant height, providing a potentially important explanation for trait variation occurring within species and independently of the environment. This work builds upon recent theoretical research developing hypotheses for how traits such as LMA should respond to increasing plant height (Falster *et al*. 2018; Westoby *et al*. 2022).

One of the key insights from our analysis was our prediction that wood density increases as plants grow taller, consistent with empirical observations based on radial wood cores (Hietz *et al*. 2013; Rungwattana & Hietz 2018). The most commonly held explanation for this phenomenon is that the mechanical stability conferred by denser wood becomes more beneficial as plants grow larger and are exposed to a higher risk of mortality due to wind stress (Hietz *et al*. 2013). Our model provides an alternative explanation based on the increasing cost of turnover and maintenance of support tissues as a fraction of net biomass production as height increases. As such, our model suggests that ontogenetic shifts in wood density may still be evolutionary advantageous even for trees recruiting in the understorey where the risk of wind exposure is relatively low (Moore *et al*. 2018).

Surprisingly, however, we observed only limited sensitivity in the response of maximum sapwood-specific conductivity to height, somewhat at odds with the strong empirical evidence for xylem conduit widening that is observed when comparing plants of increasing height at a fixed point along the stem (Olson *et al*. 2020). As described in Section 3.4, the reason that this occurred was because increasing maximum sapwood-specific conductivity also caused P_50_ to become less negative, thereby limiting the extent to which the effect of greater heights on water transport rate could be offset through variation in this trait. We note, however, that our representation of the xylem pathway was relatively simple and did not include a number of processes which may have influenced the simulated outcome such as tip-to-base widening of xylem conduits (Olson *et al*. 2021) and segmentation of vulnerability along the hydraulic pathway (Choat *et al*. 2005; Sperry & Love 2015).

### 5.4 Species turnover across soil moisture gradients and dynamic global vegetation models

Plant traits play a significant role in determining the function of terrestrial ecosystems (Westoby & Wright 2006). Consequently, trait parameterisation remains an important source of variation in the outcomes of the dynamic global vegetation models (DGVMs) (Van Bodegom *et al*. 2012) used to simulate vegetation distribution and structure and associated biogeochemical cycles in response to climate, soil, and disturbance (Prentice & Cowling 2013). Although ecosystems within DGVMs have historically been parameterised by grouping together similar species using a limited number of plant functional types (PFT), each with a prescribed set of traits, there is a growing consensus (van Bodegom *et al*. 2014; Van Bodegom *et al*. 2012; Moorcroft 2006; Sakschewski *et al*. 2015; Berzaghi *et al*. 2020; Yang *et al*. 2015; Wright *et al*. 2004) that DGVMs must move away from the PFT classification towards a continuous trait approach for two key reasons, among others. Firstly, within PFTs, there is significant variation in traits owing to species turnover and intra-specific plasticity. Secondly, *a priori* prescription of plant traits to PFTs precludes adaptation of plant strategies to future conditions (Van Bodegom *et al*. 2012).

Our exploration of species turnover across a soil moisture gradient offers a useful illustrative case for the importance of adaptation in such models. Specifically, in keeping with a productivity-drought tolerance trade-off from classical ecological theory (Smith & Huston 1989), we demonstrate how the joint optimal plant strategy conferring high productivity rates (i.e. high maximum growth rate) at the wetter end of the gradient is gradually replaced by strategies favouring drought tolerance at the drier end of the gradient (i.e. the ability to maintain positive growth rates under more negative soil water potentials). Importantly, in the absence of trait optimisation along this gradient (e.g. by following the height growth rate curve for 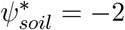 in Figure 5), we show that our growth model would underpredict the potential growth rates that could be achieved by a more optimal strategy in the wettest conditions, while simultaneously predicting non-positive growth rates under conditions which a more drought-tolerant species could persist in. Height-growth models based on the EEO framework may therefore provide a useful tool for developing the next generation of trait-based DVGMs.

### 5.5 Future directions and limitations

Our analysis based on height growth rates extends upon carbon acquisition-based EEO models by directly considering the translation of acquired carbon into future ability to acquire resources and progress towards reproductive maturity. Although height growth is likely a closer approximation of plant fitness than carbon acquisition (Potkay & Feng 2023), we note that our model omits other potentially important demographic rates which may be relevant here, including mortality risk and reproductive output. For example, taxa with greater wood density tend to have more negative lethal water potentials (Liang *et al*. 2021), and including this relationship in our model could further mediate selection towards higher wood densities in drier locations. Beyond this relationship, however, we anticipate that our findings would remain largely the same.

Perhaps the most significant opportunity for advancing the scope of the model presented here is the inclusion of a dynamic water balance and size-structured competition among individuals for water, akin to the light competition process that has been analysed in **plant** previously (Falster *et al*. 2017). Like many other EEO approaches (Wang *et al*. 2023; Dong *et al*. 2022; Xu *et al*. 2021; Trugman *et al*. 2019), trait optima emerged in our analysis for hypothetical plant individuals experiencing the environment in isolation. In reality, however, the trait optima that emerge under competition may be different due to game-theoretic interactions among strategies (Falster *et al*. 2017; Franklin *et al*. 2020). Importantly, including competition for water would permit coexistence amongst trait optima, thereby allowing inferences to made regarding the effect of water availability on functional diversity the maitenance of species diversity (Harrison *et al*. 2021; Lindh *et al*. 2014).

## Acknowledgements

Discussions with M Westoby, J Dwyer, A Pitman and I Wright enhanced the scope and quality of the analysis. IT was supported by a grant from Eucalypt Australia, AOR was supported by ARC grants to Falster & Vesk (DP200100555).

## Author contributions

IRT, PAV and DSF developed the original idea with further input from AORN and MEBS. IRT, AORN and DSF implemented development the of software required for the analysis. IRT, AORN, MEBS, PAV and DSF contributed to theoretical development of the FF16w module. IRT conducted the analysis. IRT led the writing of the manuscript, assisted by IRT, AORN, MEBS, PAV and DSF. All authors approved of the final draft of the manuscript.

## Data availability statement

The code used to conduct the analysis is available on GitHub upon request and will made available upon acceptance of the manuscript. Data are available from AusTraits (https://zenodo.org/records/3568429) and can be accessed using the AusTraits R package.

