## Supplementary Information for "Explaining plant trait variation in response to soil water availability using an optimal height-growth model"

### 6 Supporting Information

#### 6.1 Supporting Information Methods

##### 6.1.1 Photosynthesis model

Net photosynthesis, $P_{net}$, was modelled using a standard coupled stomatal-photosythnesis model, which equates photosynthetic demand for $CO_2$, $P_{net}$, with the diffusive supply of $CO_2$:

$$P_{net}(C_{\mathrm{i}}) = g_c(\psi_{\mathrm{leaf}})\frac{(C_{\mathrm{a}} - C_{\mathrm{i}})}{P_{\mathrm{atm}}}, \tag{18}$$

In turn, $P_{net}$ is modelled according to the Farquhar biochemical model (Farquhar *et al.* 1980) which includes a smoothed hyperbolic transition between two limiting rates of photosynthesis, being the Rubisco-limited rate ($A_c$) and the electron-transport limited rate ($A_j$):

$$P_{net} = \frac{A_c + A_j - \sqrt{(A_c + A_j)^2 - 4hA_cA_j}}{2h} - R_d, \tag{19}$$

where $h$ is the curvature factor of the smoothing function and $R_d$ is leaf-level respiration.

$A_c$ is defined as:

$$A_c = \frac{V_{c,max}(C_i - \Gamma^*)}{C_i + K_m}, \tag{20}$$

and $A_j$ is defined as:

$$A_j = \frac{J(C_i - \Gamma^*)}{4(C_i + 2\Gamma^*)}, \tag{21}$$

where $V_{c,max}$ is the maximum velocity of carboxylation, $C_i$ is the intercellular partial pressure of $CO_2$ and $\Gamma^*$ is the $CO_2$ compensation point.

$K_m$ is the Michaelis-Menten constant for photosythnesis and is described as:

$$K_m = K_c\left(1 + \frac{O_a}{K_o}\right), \tag{22}$$

where $K_c$ and $K_o$ are the Michaelis-Menten constants for $CO_2$ and $O_2$, respectively and $O_a$ is the atmospheric concentration of $O_2$ in terms of partial pressure.

$J$ describes the joint dependency of $A_j$ on irradiance, $I$ and the maximum electron transport rate, $J_{max}$:

$$J = \frac{aI + J_{max} - \sqrt{(aI + J_{max})^2 - 4caIJ_{max}}}{2c}, \tag{23}$$

where $a$ is the quantam yield of electron transport and $c$ is the curvature of the relationship between $J$ and $I$.

$V_{c,max}$, $J_{max}$, $\Gamma^*$, $K_c$ and $K_o$ have temperature-dependencies according to Arrhenius functions. However, because we do not consider the explicit effect of temperature in the present analysis, we assume that these parameters are fixed at their 25°C values (i.e. $V_{c,max} = V_{c,max,25}$) (Bernacchi *et al.* 2001).

##### 6.1.2 Leaf respiration

In the FF16w physiological module, leaf respiration per unit mass leaf ($r_l$; $mol\ kg^{-1}\ s^{-1}$) is assumed to be the summed component of respiration associated with nitrogen allocated to photosynthetic capacity $r_{l,p}$ and nitrogen allocated to structural components of the leaf $r_{l,s}$:

$$r_l = r_{l,s} + r_{l,p}. \tag{24}$$

Dong *et al.* (2022) show that for woody, non-nitrogen fixers, $N_{area}$ can be linearly predicted as a function of $V_{cmax,25}$ and $\phi$:

$$N_{area} = \alpha + \beta_1\phi + \beta_2 V_{c,max}. \tag{25}$$

The $\beta$ terms describes how $N_{area}$ changes with unit increases in either $\phi$ of $V_{c,max}$ and $\alpha$ is the intercept value of this equation. Estimation of $\beta_2$ from a global dataset indicates that this term corresponds with the relative mass contribution of Rubisco and cytochrome f to $V_{c,max}$ and $J_{max}$ per unit mass increase in $V_{c,max}$, assuming as we do here, that $V_{c,max,25}$ and $J_{max}$ occur in the leaf at a ratio of approximately 2:1 (Smith & Keenan 2020; Dong *et al.* 2022).

Assuming that $\alpha$ accounts for structural components of $N_{area}$ not accounted for by $\phi$, it can be argued that:

$$N_{mass,s} = (\alpha + \beta_1 \phi)\phi^{-1}, \tag{26}$$

and that,

$$N_{mass,p} = (\beta_2 V_{c,max})\phi^{-1}. \tag{27}$$

Then, $N_{mass}$ is multiplied by $R$, the rate of respiration per unit mass nitrogen to give:

$$r_{l,s} = R_{l,s} N_{mass,s}, \tag{28}$$

and,

$$r_{l,p} = R_{l,p} N_{mass,p}. \tag{29}$$

Importantly, $R_{l,s}$ and $R_{l,p}$ can be different values, reflecting possible differentiation in the repspiration rate of a given mass of structural versus photosynthetic nitrogen.

#### 6.2 Supporting Information Tables

Table S1: **Variable descriptions, tested parameter values and units in the Supplementary Information Methods**.

| Symbol | Description | Tested values | Units |
|---|---|---|---|
| **Parameter** | | | |
| $K_{c,25}$ | Michaelis-Menten constant (carboxylation) | 404.9 | $\mu$mol mol$^{-1}$ |
| $K_{o,25}$ | Michaelis-Menten constant (oxygenation) | 278.4 | mmol mol$^{-1}$ |
| $\Gamma^*_{25}$ | $CO_2$ compensation point | 42.75 | $\mu$mol mol$^{-1}$ |
| $h$ | Curvature factor | 0.99 | unitless |
| $N_{mass,s}$ | Structural nitrogen mass per leaf mass | Derived from $\phi$ | kg N kg$^{-1}$ |
| $N_{mass,p}$ | Photosynthetic nitrogen mass per leaf mass | Derived from $\phi$, $V_{c,max}$ | kg N kg$^{-1}$ |
| $R_{l,p}$ | Respiration per unit mass photosynthetic nitrogen | 40000 | mol yr$^{-1}$ kg N$^{-1}$ |
| $R_{l,s}$ | Respiration per unit mass structural nitrogen | 21000 | mol yr$^{-1}$ kg N$^{-1}$ |
| $r_{l,p}$ | Respiration per unit mass leaf associated with photosynthetic nitrogen | Derived from $N_{mass,p}$, $R_{l,p}$ | mol yr$^{-1}$ kg$^{-1}$ |
| $r_{l,s}$ | Respiration per unit mass leaf associated with structural nitrogen | Derived from $N_{mass,s}$, $R_{l,p}$ | mol yr$^{-1}$ kg$^{-1}$ |
| $\alpha$ | Intercept of $N_{area}$ empirical relationship | 0.535[a] | unitless |
| $\beta_1$ | Linear coefficient for $\phi$ in $N_{area}$ empirical relationship | 0.009[a] | unitless |
| $\beta_2$ | Linear coefficient for $V_{c,max}$ in $N_{area}$ empirical relationship | 0.006[a] | unitless |

[a] Dong *et al.* (2022)

##### 6.3 Supporting Information Figures

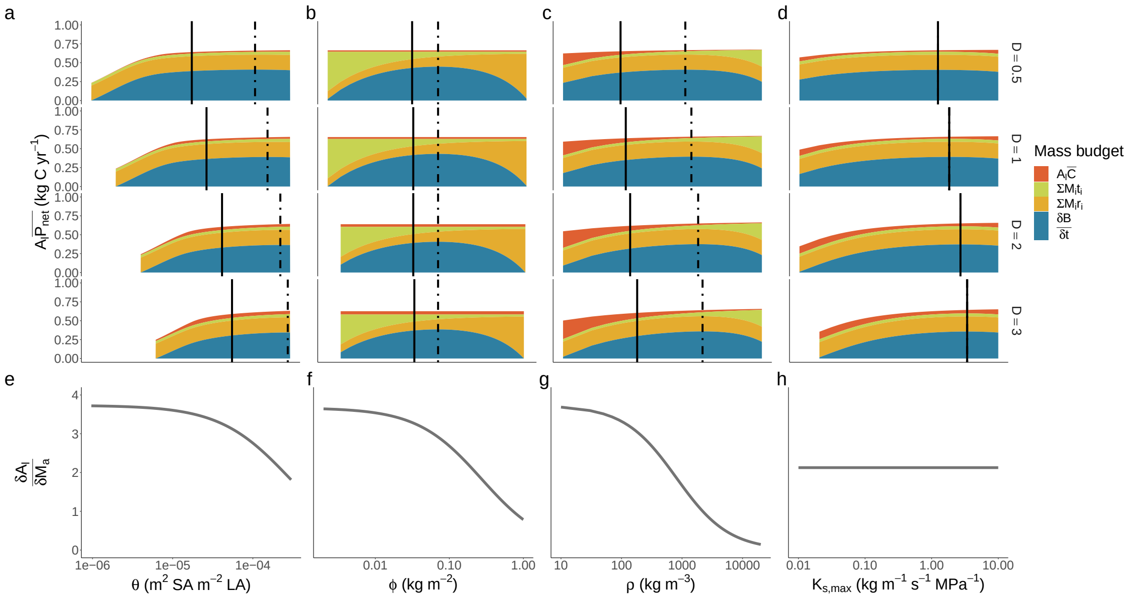


Figure S1: **Decomposition of the components determining trait optima across a vapour pressure deficit (D) gradient**. The top row of panels shows how net biomass production, $\frac{\mathrm{d}B}{\mathrm{d}t}$ emerges for each trait across an increasing vapour pressure deficit gradient (i.e. from the top to the bottom of panels **a-d**) as the residual of total assimilation, $(A_l \bar{P_{net}})$, after accounting for hydraulic costs, $A_l\bar{C}$, turnover $\sum M_i t_i$ and respiration $\sum M_i r_i$ of each plant tissue, $i$. The trait value maximising $\frac{\mathrm{d}B}{\mathrm{d}t}$ is indicated by the dashed vertical bar. The solid vertical line indicates the trait value maximising the height-growth rate, $\frac{dH}{dt}$. The $\frac{dH}{dt}$ optima emerges through multiplication of $\frac{dB}{dt}$ with the rate of leaf area deployment per unit of live mass growth $\frac{\mathrm{d}A_l}{\mathrm{d}M_a}$, shown in the second column. For most traits, this causes the $\frac{dH}{dt}$ optima to be lower than the $\frac{dB}{dt}$ optima, owing to the greater value of $\frac{\mathrm{d}A_l}{\mathrm{d}M_a}$ at low trait values in panels **e-g** but also explains why the optima are equivalent for $K_{s,max}$.

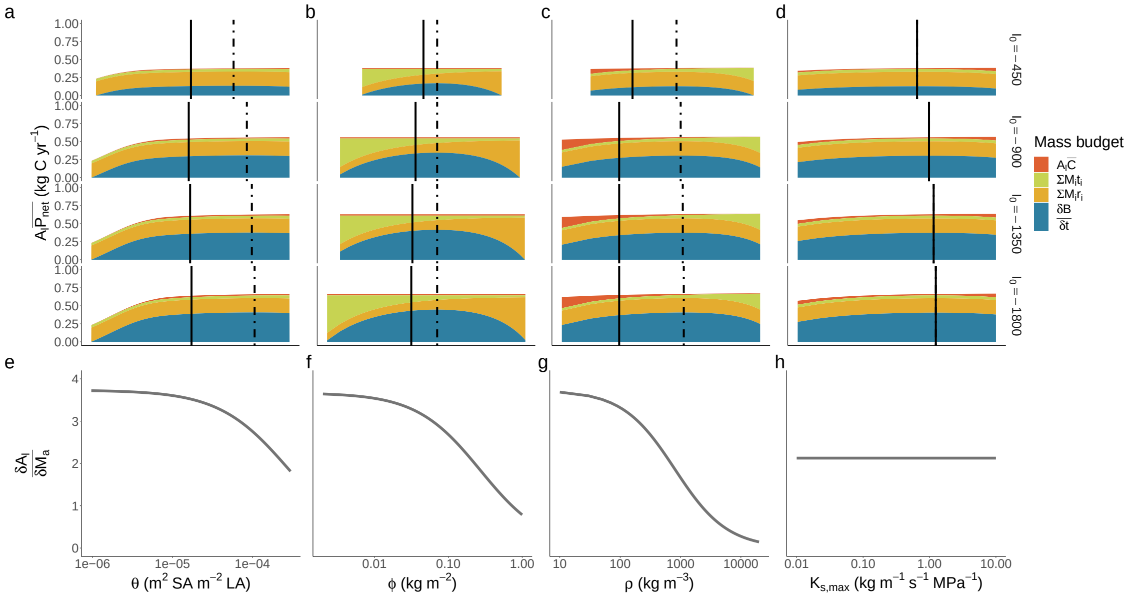


Figure S2: **Decomposition of the components determining trait optima across a above-canopy photon plux density ($I_0$) gradient**. The top row of panels shows how net biomass production, $\frac{\mathrm{d}B}{\mathrm{d}t}$ emerges for each trait across a increasing light availability gradient (i.e. from the top to the bottom of panels **a-d**) as the residual of total assimilation, $(A_l \bar{P_{net}})$, after accounting for hydraulic costs, $A_l\bar{C}$, turnover $\sum M_i t_i$ and respiration $\sum M_i r_i$ of each plant tissue, $i$. The trait value maximising $\frac{\mathrm{d}B}{\mathrm{d}t}$ is indicated by the dashed vertical bar. The solid vertical line indicates the trait value maximising the height-growth rate, $\frac{dH}{dt}$. The $\frac{dH}{dt}$ optima emerges through multiplication of $\frac{dB}{dt}$ with the rate of leaf area deployment per unit of live mass growth $\frac{\mathrm{d}A_l}{\mathrm{d}M_a}$, shown in the second column. For most traits, this causes the $\frac{dH}{dt}$ optima to be lower than the $\frac{dB}{dt}$ optima, owing to the greater value of $\frac{\mathrm{d}A_l}{\mathrm{d}M_a}$ at low trait values in panels **e-g** but also explains why the optima are equivalent for $K_{s,max}$.

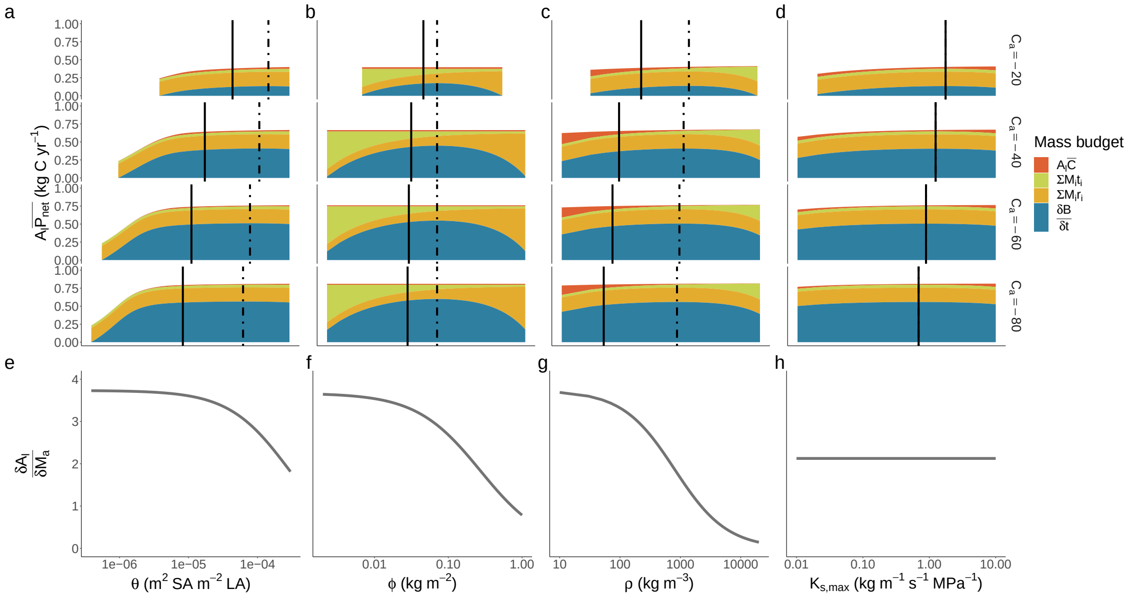


Figure S3: **Decomposition of the components determining trait optima across a atmospheric $CO_2$ gradient** ($C_a$). The top row of panels shows how net biomass production, $\frac{dB}{dt}$ emerges for each trait across an increasing atmospheric concentration of $CO_2$ (i.e. from the top to the bottom of panels **a-d**) as the residual of total assimilation, $(A_l\bar{P_{net}})$, after accounting for hydraulic costs, $A_l\bar{C}$, turnover $\sum M_i t_i$ and respiration $\sum M_i r_i$ of each plant tissue, $i$. The trait value maximising $\frac{dB}{dt}$ is indicated by the dashed vertical bar. The solid vertical line indicates the trait value maximising the height-growth rate, $\frac{dH}{dt}$. The $\frac{dH}{dt}$ optima emerges through multiplication of $\frac{dB}{dt}$ with the rate of leaf area deployment per unit of live mass growth $\frac{dA_l}{dM_a}$, shown in the second column. For most traits, this causes the $\frac{dH}{dt}$ optima to be lower than the $\frac{dB}{dt}$ optima, owing to the greater value of $\frac{dA_l}{dM_a}$ at low trait values in panels **e-g** but also explains why the optima are equivalent for $K_{s,max}$.

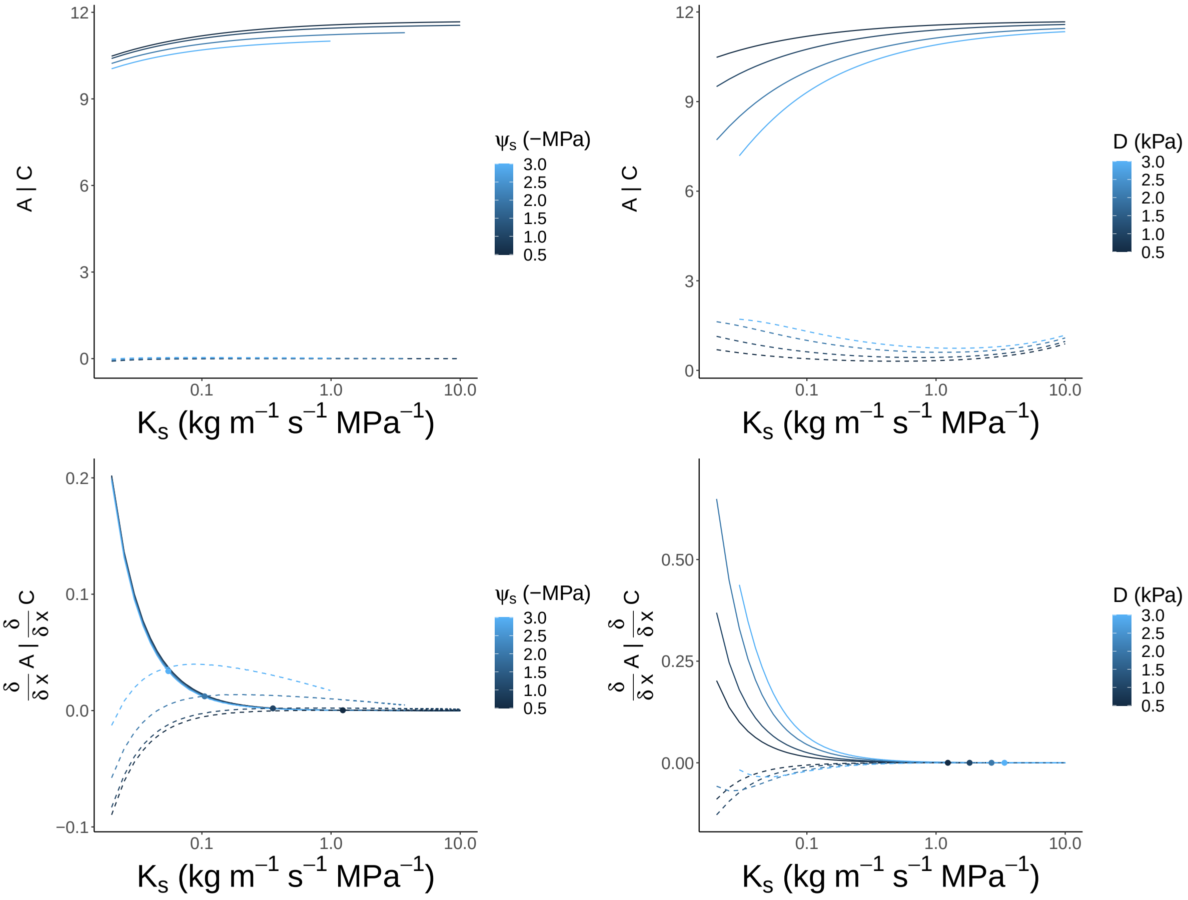


Figure S4: Graphical explanation for the response of $K_{s,max}$ to soil water availability ($\psi_{soil}$ and atmospheric vapour pressure deficit (D). In the top row of plots, the optimal photosynthetic assimilation (A; solid lines) and hydraulic costs (C; dotted lines) from the stomatal model for a given set of traits (x) are plotted with respect to variation in $K_{s,max}$. Colour shading indicates the environmental conditions that the stomatal model was optimised under. In the bottom row of plots, the same curves derived with respect to variation in $K_{s,max}$ are illustrated. The optimum $K_{s,max}$ under each environment is indicated by the coloured points and occurs where $\frac{\delta}{\delta x}A = \frac{\delta}{\delta x}C$ The left column of plots illustrates how $K_{s,max}$ increases as soils dry owing to an increase in the elevation of $\frac{\delta}{\delta x}C$. The right column of plots illustrates how $K_{s,max}$ increases with D owing primarily to an increase in the elevation of $\frac{\delta}{\delta x}A$, which occurs because $A$ increases more rapidly with $K_{s,max}$ in drier atmospheres.

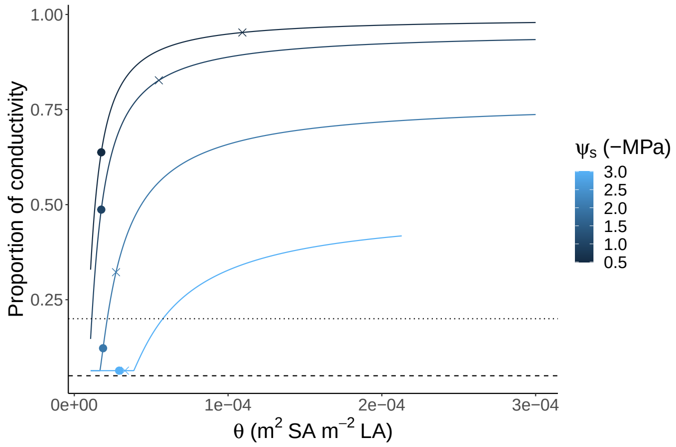


Figure S5: The proportion of conductivity in the xylem increases with the sapwood to leaf area ratio ($\theta$) because $\psi_{\mathrm{leaf}}$ is less negative when the water transport rate is higher. As $\psi_{\mathrm{soil}}$ becomes more negative, the biomass production (crosses) and height-growth rate (circles) optimising values of $\theta$ decline, because the costs associated with greater sapwood are greater than the cost of losing hydraulic conductivity, except in very dry soils (lightest blue). The horizontal lines indicate losses of conductivity associated with a high risk of drought-based mortality (80%; Hammond *et al.* 2019) and catastrophic xylem failure (95%).
